# EZH2 endorses cell plasticity to carcinoma cells facilitating mesenchymal to epithelial transition and tumour colonization

**DOI:** 10.1101/2022.03.11.483977

**Authors:** Amador Gallardo, Aldara Molina, Helena G Asenjo, Lourdes Lopez-Onieva, Juan Carlos Alvarez-Perez, Jordi Martorell-Marugán, Francisca E Cara, Mencia Espinosa-Martinez, Pedro Carmona-Sáez, Sergio Granados-Principal, Pedro P Medina, Antonio Sanchez-Pozo, David Landeira

## Abstract

Reversible transition between the epithelial and mesenchymal states are key aspects of carcinoma cell dissemination and the metastatic disease, and thus, characterizing the molecular basis of the epithelial to mesenchymal transition (EMT) is crucial to find druggable targets and more effective therapeutic approaches in cancer. Emerging studies suggest that epigenetic regulators might endorse cancer cells with the cell plasticity required to conduct dynamic changes in cell state during EMT. However, epigenetic mechanisms involved remain mostly unknown. Polycomb Repressive Complexes (PRCs) proteins are well-established epigenetic regulators of development and stem cell differentiation, but their role in different cancer systems is inconsistent and sometimes paradoxical. In this study, we have analysed the role of the PRC2 protein EZH2 in lung carcinoma cells. We found that besides its described role in *CDKN2A*-dependent cell proliferation, EZH2 upholds the epithelial state of cancer cells by repressing the transcription of hundreds of mesenchymal genes. Chemical inhibition or genetic removal of EZH2 promotes the residence of cancer cells in the mesenchymal state during reversible epithelial-mesenchymal transition. In fitting, analysis of human patient samples and tumour xenograft models indicate that EZH2 is required to efficiently repress mesenchymal genes and facilitate tumour colonization *in vivo*. Overall, this study discloses a novel role of PRC2 as a master regulator of EMT in carcinoma cells. This finding has important implications for the design of therapies based on EZH2 inhibitors in human cancer patients.

## Introduction

Epithelial to mesenchymal transition (EMT) is a cellular program crucial for embryogenesis, wound healing and malignant progression. EMT mediates dispersion of cells in embryos, formation of mesenchymal cells in injured tissues and initiation of the invasive and metastatic behaviour of epithelial cancers ^1,2^. EMT is a dynamic and reversible process between the epithelial and the mesenchymal cell states, and therefore, different types of metastable quasimesenchymal cells can revert back to an epithelial state in a process known as mesenchymal– epithelial transition (MET) ^3,4^. The current view is that EMT is governed by the activity of the EMT-inducing transcription factors (EMT-TFs), TWIST1/2, SNAI1/2 and ZEB1/2, that coordinate the repression of genes that maintain the epithelial state and the activation of the ones that induce the acquisition of mesenchymal features ^3,5^. Expression of EMT-TFs is co-ordinately regulated by several intracellular signalling pathways of which the most prominent is the transforming growth factor-β (TGF-β) pathway that can on its own induce entrance into the EMT program ^6^. The molecular mechanisms facilitating MET remain mostly unknown. Although EMT-MET is in essence a dynamic change between cellular states that must be regulated at the epigenetic level, chromatin modifiers involved and their mode of action remain to be identified and characterized ^3,4^. In the context of cancer, activation of EMT imparts features that are essential to the formation of metastasis by carcinoma cells, including cell motility, ability to disseminate, tumour-initiating properties and elevated resistance to chemotherapy ^7,8^. In addition, MET is associated with more efficient colonization of new tissues by carcinoma cells at metastatic sites ^3,7^. Thus, characterizing the molecular basis of EMT and MET is crucial to better understand cancer progression and design more precise therapeutic intervention approaches.

Polycomb group (PcG) proteins are chromatin regulators of human development, tissue homeostasis and cancer ^9,10^. PcG proteins associate to form multimeric complexes termed Polycomb Repressive Complex 1 and 2 (PRC1 and PRC2) of which the catalytic subunits are RING1A/B and EZH1/2 respectively. In stem cells, RING1A/B monoubiquitinates lysine 119 on histone H2A (H2AK119ub) while EZH1/2 tri-methylates lysine 27 on histone H3 (H3K27me3) coordinately to establish chromatin domains that maintain transcriptional repression of hundreds of lineage-specific genes, reenforcing maintenance of the current gene expression program and upholding cell identity ^9,10^. Importantly, although mutations of genes that encode for PcG proteins are recurrent and associated with poor prognosis in a large number of human cancers, the role of PRCs during malignant progression remains unclear ^10,11^. Initial studies showed that PcG proteins acted as oncogenes by repressing the tumour suppressor *INK4A/ARF* (*CDKN2A*) locus, and therefore inhibiting senescence and favouring cell proliferation ^12,13^. However, subsequent studies revealed senescence-independent prooncogenic as well as tumour suppressor functions, highlighting unanticipated complexity in the role of PcG proteins in cancer ^11,14^. This apparently conflicting observations are probably a consequence of the ability of PRC2 to regulate hundreds of functionally distinct target genes, together with the fact that PRC2 function during cancer progression can be dramatically affected by the genetic alterations accumulated by the cancer cell ^15–18^.

The initial oncogenic role of EZH2 led to the development of several small molecule inhibitors that are currently being tested in clinical trials ^19^. Thus, given the context-dependent function of EZH2 in cancer, it is urgent a much better understanding of its molecular role in this disease. Emerging studies have led to hypothesize that the function of PcG proteins during cancer progression does not only rely on the regulation of cell proliferation through the *CDKN2A* locus, but also on the ability of PRC2 to maintain the repression of hundreds of genes in the tumour cell of origin, which in turn might regulate epigenetic plasticity and impact cancer dissemination ^11,20^. In fitting with this idea, PRC2 can affect changes in cell state during EMT, but current studies reveal apparently contradictory results lacking a unifying molecular mechanism ^18,21,22^.

Here, we set out to determine the molecular mechanism by which PRC2 regulates EMT-MET in human lung carcinoma cells. We established a TGF-β-inducible reversible *in vitro* system and we found that EZH2 facilitates the maintenance of the epithelial identity by binding and repressing the transcription of a large set of mesenchymal genes during EMT-MET. These include well-known regulators of the mesenchymal state such as *SNAI2, MMP2* and *ITGB3*. In agreement, analysis of human patient samples and xenograft experiments support that EZH2 is required to efficiently repress mesenchymal genes and facilitate tumour colonization *in vivo*. Overall, our results support that EZH2 is required to repress mesenchymal genes during MET and facilitate tumour colonization in lung cancer.

## Results

### EZH2 binds and represses mesenchymal genes in lung carcinoma cells

To dissect the function of EZH2 in lung cancer we first focused on the human A549 cell line, which is a well-established system to study the molecular basis of non-small cell lung cancer. This cell line is homozygous null for *CDKN2A* and thus it is suitable to study the *CDKN2A*-independent role of EZH2 in cancer. In addition, A549 cells harbour a homozygous point mutation in the *KRAS* gene (G12S) encoding a hyperactive isoform of KRAS protein. First, we analysed the genome-wide binding profile of EZH2 by chromatin immunoprecipitation followed by high throughput sequencing (ChIP-seq). We found that, in agreement with the role of this protein in stem cell biology, there were 1969 EZH2 binding peaks that mapped to the promoter of 1237 genes (Figure 1A, S1A), suggesting that PRC2 is repressing transcription of this set of genes. In pluripotent stem cells, promoters bound by EZH2 display bivalent chromatin and thus they are not only enriched for H3K27me3 but are also marked by H3K4me3 ^23,24^. Likewise, analysis of publicly available H3K27me3 and H3K4me3 ChIP-seq datasets confirmed that most of EZH2 target promoters in A549 cells displayed a bivalent state (Figure 1A, B, S1B), and that there is a positive correlation between the level of EZH2 and H3K27me3 at individual target promoters (Figure 1C). As expected, transcriptome profiling of A549 cells by mRNA-high throughput sequencing (mRNA-seq) showed that EZH2 target genes were transcriptionally repressed or expressed at very low level (Figure 1D). EZH2-repressed bivalent domains were readily detectable at individual target regions (i.e. *GREM1* gene in Figure 1E). Importantly, gene ontology analysis revealed that many EZH2 target genes are related to mesenchymal cells (Figure 1F). Thus, similarly to the role of EZH2 in stem cells, we concluded that EZH2 binds to hundreds of bivalent genes that are transcriptionally repressed in A549 cells. Interestingly, EZH2-target genes were enriched in mesenchyme-related terms leading us to hypothesize that EZH2 is required to repress the mesenchymal gene expression program and maintain an epithelial cell state in A549 cells.

**Figure 1.**
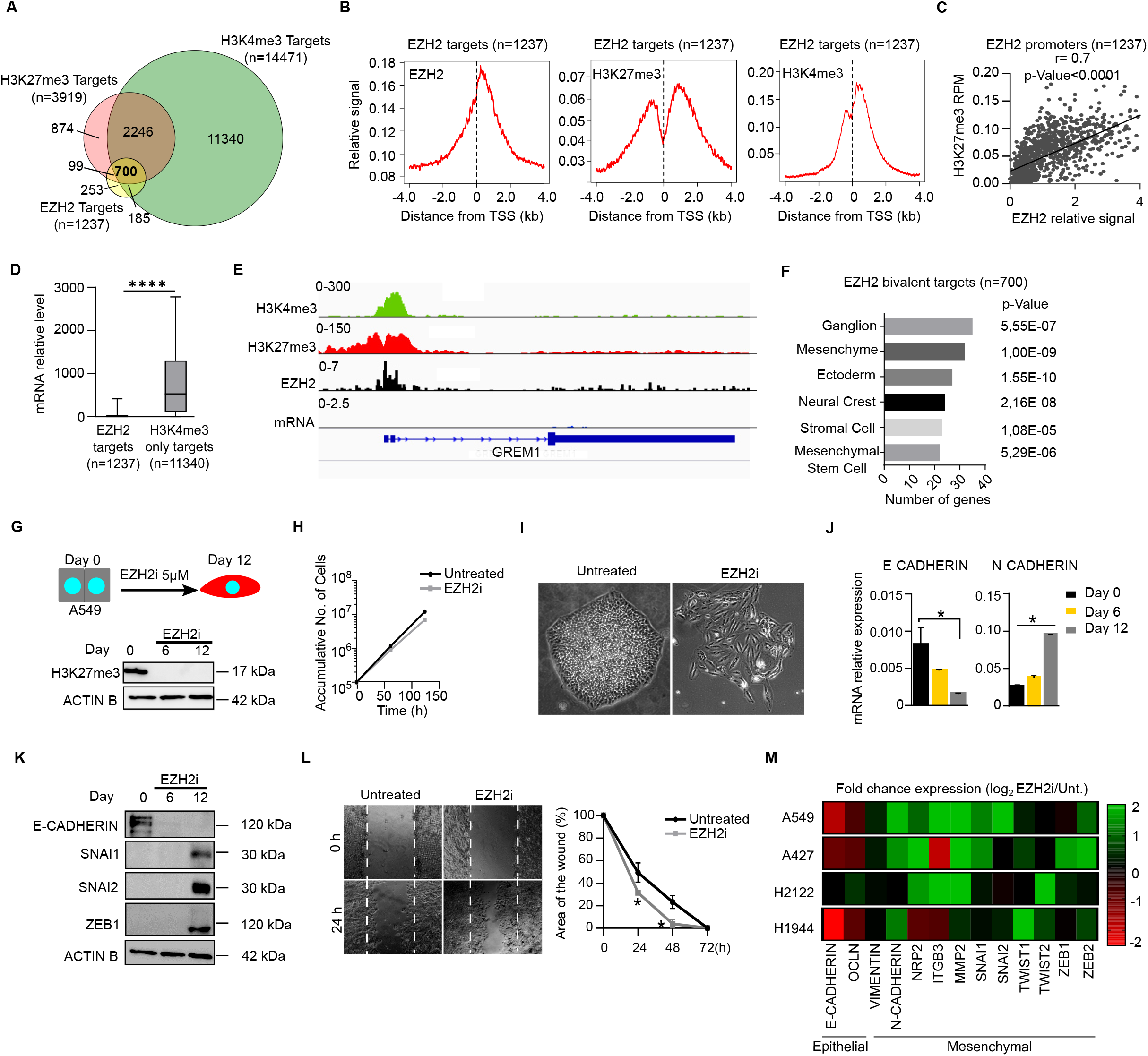
EZH2 binds and represses mesenchymal genes in lung carcinoma cells. **(A)** Venn diagram comparing gene promoters bound by EZH2, H3K27me3 and H3K4me3 in ChIP-seq analyses in A549 cells. **(B)** ChIP-seq average binding profile of EZH2, H3K27me3 and H3K4me3 around the TSS of 1237 gene promoters bound by EZH2 in A549 cells. **(C)** Spearman correlation analysis of the binding of EZH2 and H3K27me3 at EZH2-target genes. **(D)** Boxplot of mRNA transcribed from EZH2-target or control promoters measure by RNA-seq. **(E)** Genome browser view of H3K4me3, H3K27me3, EZH2 binding and mRNA level at the *GREM1* locus. **(F)** Gene Ontology analysis using Jensen tissues database of EZH2-bound bivalent targets. **(G)** Schematic diagram of the treatment of A549 cells with 5 μM EZH2i and western blot analysis of whole-cell extracts comparing the level of H3K27me3 during the course of the experiment. ACTIN B provides a loading control. **(H)** Growth curve of A549 cells untreated or treated with EZH2i. **(I)** Brightfield images of A549 cells plated at low density and treated with EZH2i or untreated during 14 days. **(J)** Analysis of *E-CADHERIN* and *N-CADHERIN* mRNA expression by RT-qPCR during EZH2i treatment in A549 cells. Expression is calculated relative to housekeeping genes *GAPDH* and *ACTINB*. **(K)** Western blot analysis of whole-cell extracts in EZH2i-treated A549 cells. ACTIN B was used as a loading control. **(L)** Brightfield images and plot comparing A549 cells untreated or pretreated with EZH2i for four days and plated in the presence of EZH2i during the course of a wound healing assay. **(M)** Heatmap representing the fold change in the expression of epithelial and mesenchymal markers (treated vs untreated with EZH2i) in the indicated NSCLC cell lines (A549, A427, H2122, H1944). Mean and SEM of 3 experiments are shown in H, J and L. Asterisks indicate statistical significance using a Mann-Whitney test (* p<0.05, **** p<0.0001).

To address whether EZH2 was upholding the epithelial state in A549 cells we analysed the effect of inhibiting EZH2 methyltransferase activity. We used a highly specific small molecule inhibitor of EZH2 (GSK126/EZH2i) ^25^. Titration experiments confirmed that 5 μM EZH2i drastically reduces global levels of H3K27me3 upon four days of treatment, leaving only a residual H3K27me3 signal that is probably a consequence of EZH1 activity (Figure S1C). Importantly, growing A549 cells in the presence of EZH2i for twelve days, consistently reduced global H3K27me3 levels without affecting cell growth (Figure 1G, H). Strikingly, A549 cells plated at low density and grown in the presence of the EZH2i for 18 days displayed a very clear phenotype as compared to untreated cells: they formed unpacked colonies with more elongated and fusiform cell morphology typically associated with mesenchymal states (Figure 1I, S1D). In fitting, EZH2i-treated cells displayed reduced levels of the epithelial marker *E-CADHERIN* and augmented expression of the mesenchymal markers such as *N-CADHERIN* and EMT-TFs including SNAI1, SNAI2 and ZEB1 (Figure 1J, K, S1E). Among the analysed EMT-TFs, upregulation of SNAI2 protein was associated with increased abundance of its mRNA transcript, suggesting that EZH2 directly represses transcription of *SNAI2* (Figure S1E). In fitting, ChIP-qPCR analysis confirmed that SNAI2 is a direct target of EZH2 activity (Figure S1F).

Next, we asked whether in addition to the upregulation of mesenchymal molecular markers, treatment with EZH2i led to acquisition of mesenchymal cellular features. In agreement with our molecular analyses, cultured wound healing assays showed that EZH2i-treated cells healed faster that control cells (Figure 1L). Complementarily, inhibition of EZH2 resulted in cells with reduced capacity to grown as colonies in soft agar (Figure S1G) and as tumourspheres (Figure S1H). Thus, we concluded that inhibition of EZH2 in A549 cells leads to the expression of EMT-TFs and acquisition of mesenchymal cellular features.

To confirm that these findings were not exclusive feature of the A549 cell line, we analysed whether inhibition of EZH2 induced activation of mesenchymal genes in three additional lung cancer cell lines that are genetic null for *CDKN2A* (cell lines A427, H1944 and H2122). In fitting with our results in A549 cells, treatment with 5 μM EZH2i during eight days had only minor effects on cell survival in all tested cell lines (Figure S1I). Importantly, plating cells at low density in the presence of EZH2i promoted the upregulation of mesenchymal genes (Figure 1M) and the acquisition of a typically mesenchymal cellular morphology in the four tested cell lines; unpacked colonies composed by fusiform cells (Figure S1J). We concluded that EZH2 binds and represses mesenchymal genes, and that inhibition of EZH2 promotes spontaneous transition into a quasi-mesenchymal state in NSCLC cell lines.

### TGF-β-induced EMT is reversible in A549 cells

To analyse the function of EZH2 during EMT in lung cancer we setup an *in vitro* reversible system to dissect the molecular mechanisms regulating EMT and MET. We titrated TGF-β and epidermal growth factor (EGF) and determine that treatment of A549 cells with 10 ng/ml TGF-β and 50 ng/ml EGF rapidly induced downregulation of E-CADHERIN mRNA and increased wound-healing capacity after 48 hours of treatment in A549 cells (Figure S2A, B). Treatment of cells with TGF-β + EGF during six days induced clear morphological changes in cell culture: cells stopped forming tight colonies and acquired a fusiform shape (Day6, Mesenchymal (M)) (Figure 2A). Removal of both cytokines from the culture media prompted a reversion of cell morphology to the epithelial state within the following six days (Day 12, epithelial reverted (ER)) (Figure 2A). In agreement, time course analysis of the expression of epithelial (*E-CADHERIN*) and mesenchymal (*N-CADHERIN*, *NRP2*) markers showed expected opposite patterns of expression (Figure 2B, C). To address whether the presence or absence of TGF-β + EGF in the culture media led to extensive reorganization of epithelial-mesenchymal gene expression programs, we compared the transcriptome of cells at days 0, 6 and 12 by mRNA sequencing (mRNA-seq). Cells at day 0 and 6 differentially expressed 1485 genes of which most of them (936 genes) were upregulated upon EMT on day 6 (Figure S2C, D). Gene ontology analysis confirmed that these genes were enriched in mesenchymal-related terms including mesenchymal tissues (stromal cells, mesenchymal stem cells), cellular processes (focal adhesion, cell migration) and signalling pathways (Ras and PI3K-Akt signalling) (Figure S2E). As expected, comparison of cells in days 6 and 12 showed the opposite trend: most differentially expressed genes were downregulated (586 out of 990 genes) and these were again associated with a mesenchymal phenotype (Figure S2F, G, H). Importantly, a large set of the differentially expressed genes were reversibly regulated: they were either downregulated or upregulated on day 6, but restored to their initial expression level on day 12 (829 reversible genes, Figure 2D, Table S1). Reversible genes upregulated on day 6 were again enriched in ontology terms associated with mesenchyme (Cluster II, Figure 2D) and included key genes with important roles in a wide range of cellular processes (i.e. cell surface, ligands, extracellular matrix organization and transcription factors) (Figure 2E). Analysis of key rationally-selected epithelial (i.e. *E-CADHERIN, CLDN2, OCLN*) and mesenchymal markers (*N-CADHERIN, VIMENTIN, MMP2/9/15/25*, EMT-TFs) also displayed a reversible gene expression pattern during the course of the experiment (Figure 2F, G). In fitting, analysis of protein expression by Western-blot confirmed that the level of EMT-TFs (SNAI1/2 and ZEB1) peaked on day 6 and was reduced on day 12 (Figure 2H). Importantly, comparison of wound-healing capacity at different time points showed that M cells on day 6 heal faster than E and ER cells on day 0 and 12 (Figure 2I). Taken together, these analyses indicate that addition of TGF-β and EGF to the culture media induces the activation of a mesenchymal gene expression program and mesenchymal cellular features, and that removal of these cytokines facilitates the reversion of the cells to an epithelial state. We concluded that transient addition of TGF-β and EGF to A549 cells provides a well-suited system to analyse the molecular basis of EMT-MET in lung cancer.

**Figure 2.**
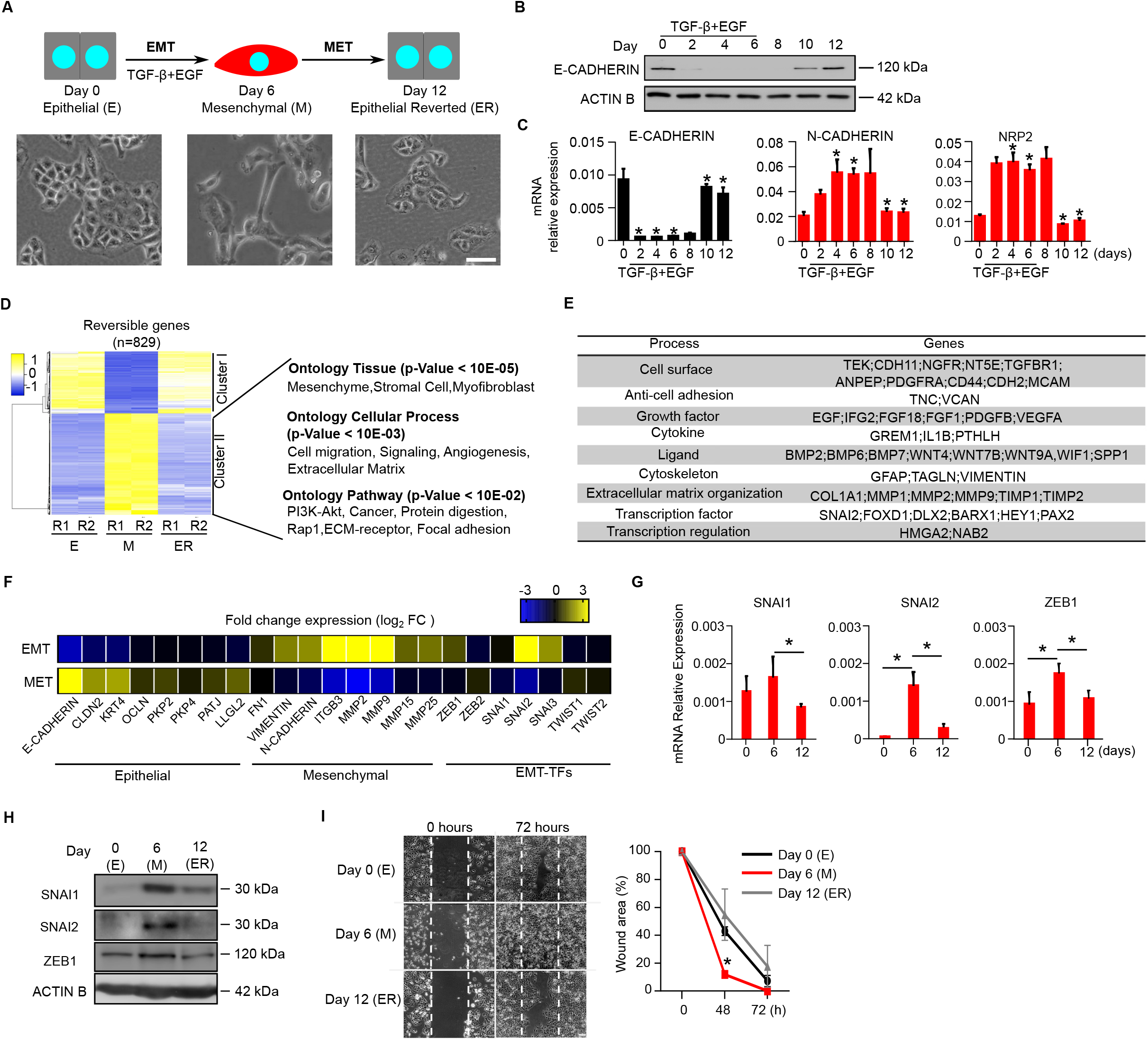
TGF-β-induced EMT is reversible in A549 cells. **(A)** Scheme of the experimental design used to induce EMT and MET in A549 cells during 12 days. Brightfield images of the representative morphology at the different stages of the transitions are shown. Scale bar 100uM. **(B)** Expression of *E-CADHERIN* by western-blot during the course of the experiment. ACTIN B serves as loading control. **(C)** Analysis of mRNA expression by RT-qPCR of indicated epithelial or mesenchymal genes during the course of the experiment. Relative expression level against *GAPDH* and *ACTIN B* is shown. **(D)** Heatmap analysis of mRNA expression of 829 reversibly regulated genes (FC>2, p<0.05) at day 0 (E), day 6 (M) and day 12 (ER) by RNA-seq of two independent replicates (R1 and R2). Yellow and blue indicate higher and lower expression respectively. Gene ontology analyses of genes in cluster II are shown. **(E)** Table showing reversible genes included in the GO category “Mesenchyme” identified in (D). **(F)** Heatmaps representing the fold change expression measured by mRNA-seq of rationally-selected epithelial and mesenchymal markers during EMT (upper strip) or MET (lower strip) **(G)** Analysis of mRNA expression by RT-qPCR of indicated EMT-TFs during EMT-MET. Expression level relative to *GAPDH* and *ACTIN B* is shown. **(H)** Analysis of the expression of SNAI1, SNAI2 and ZEB1 proteins during EMT-MET by Western blot. ACTIN B was used as a loading control. **(I)** Brightfield images and quantification plot comparing the wound healing capacity of cells at day 0, 6 and 12 of EMT-MET. Mean and SEM of 3 experiments are shown in B, G and I. Asterisks indicate statistical significance using a Mann-Whitney test (* p<0.05).

### EZH2 directly regulates a large set of mesenchymal genes during EMT-MET in lung cancer cells

To dissect the role of EZH2 during epithelial-mesenchymal transition we first focused our analysis in the expression level of PRC2 components during EMT. EZH2, other PRC2 subunits and H3K27me3 were detected at similar levels at days 0 (E) and day 6 (M) (Figure 3A, B, C, S3A, B), suggesting that PRC2 is functional in both epithelial and mesenchymal cell states. Quantitative ChIP-seq (cChIP-seq) analysis of EZH2 binding at day 0 (E) and day 6 (M) revealed that EZH2 can bind to 1237 gene promoters in epithelial and/or mesenchymal states (Table S1). Strikingly, despite widespread changes in gene expression were detected upon EMT (Figure 2D), EZH2 displayed only minor changes in the pattern of binding of EZH2 around the transcription start site (TSS) of target genes in both cell states (Figure 3D, E, S3C). To confirm this observation, we focused our analysis on genes that were bound by EZH2 and were transcriptionally up-regulated (FC>2, p<0.05) during EMT (386 genes, Figure 3F, Table S1). Albeit these genes showed clear binding of EZH2, H3K27me3 and H3K4me3 around the TSS at day 0 (Figure 3G, S3D), and were transcriptionally induced on day 6 (Figure 3H), binding of EZH2 to the promoter region remained mostly unaltered during the six days of EMT induction (Figure 3I). This suggests that although EZH2 is repressing mesenchymal target genes in the epithelial state, their transcriptional upregulation during EMT does not require eviction of EZH2 from target promoters (i.e. *BMP2*, Figure 3J). A change in the binding of EZH2 was detected for a few genomic regions (i.e. *RNF182*) (Figure S3E), demonstrating that stable binding of EZH2 to target promoters is not a technical caveat of our cChIP-seq approach. In fitting, stable binding of EZH2 to target gene promoters (*FGF3*, *LGR5*, *NPTX1* and *CCND2*) was also detected by ChIP-PCR (Figure S3F). Previous reports show that PRC2 represses the *E-CADHERIN* gene during EMT in prostate and colon cancer cell lines (Cao et al. 2008; Herranz et al. 2008). However, we only found a marginal increased binding of EZH2 to the promoter region of E-cadherin on day 6 in A549 cells (Figure S3F, G). Thus, we concluded that EZH2 binds to 386 mesenchymal genes that are transcriptionally induced upon TGF-β-stimulation, and that their activation does not require eviction of EZH2 from target promoters.

**Figure 3.**
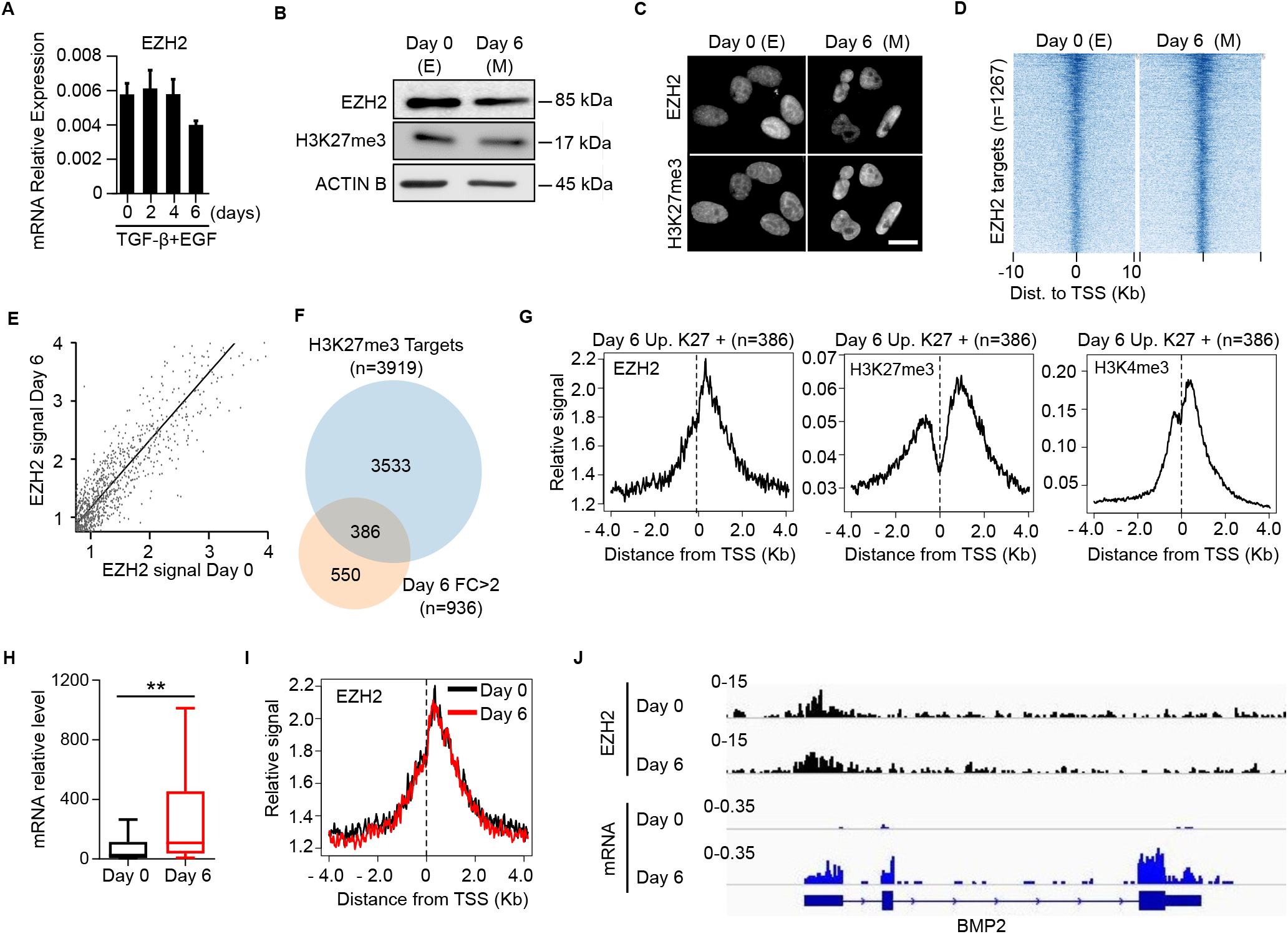
Binding of EZH2 to target promoters remains constant during EMT in A549 cells. **(A)** Expression of EZH2 mRNA by RT-PCR at different times points during EMT. **(B)** Western-blot analysis at day 0 (E) and day 6 (M) during EMT. ACTIN B was used as loading control. **(C)** Immunofluorescence analyses of EZH2 and H3K27me3 at day 0 (E) and day 6 **(M)** during EMT. Scale bar is 10 μm. **(D)** Heatmap of the EZH2 cChIP-seq signal around the TSS of 1267 genes bound by EZH2 in cells at day 0 (E) and/or day 6 (M). **(E)** Correlation analysis of EZH2 signal at target promoters (−0.5+1.5Kb around the TSS) in cells at day 0 (E) and/or day 6 (M). **(F)** Venn diagram comparing genes marked by H3K27me3 at day 0 (E) and up-regulated (FC>2, p<0.05) on day 6 (M). **(G)** Average binding profile of indicated proteins at 386 gene promoters identified in F. **(H)** Analysis of mRNA expression from 386 gene promoters identified in F at day 0 (E) and day 6 (M). Asterisks indicate statistical significance using Mann-Whitney test (**p<0.01). **(I)** Average binding profile of EZH2 at day 0 (E) and day 6 (M) at 386 gene promoters identified in (F). **(J)** Genome browser view of EZH2 binding and mRNA expression at the *BMP2* locus at day 0 (E) and day 6 (M).

To address how the loss of function of EZH2 affects the regulation of its target genes during EMT-MET, we compared the transcriptome of EZH2i-treated and control cells during EMT-MET (Figure 4A). A549 cells were pre-treated with 5 μm EZH2i for four days to reduce H3K27me3 to background levels and EZH2i was refreshed every two days during the course of the experiment, leading to sustained reduction of H3K27me3 levels (Figure S4A). TGF-β-treated cells acquired mesenchymal morphology on day 6, irrespectively of being treated or not with EZH2i (Figure S4B). In contrast, while untreated cells reverted back to an epithelial morphology on day 12, EZH2i-treated cells retained a clear mesenchymal morphology (Figure S4B). In agreement, examination of the expression of key epithelial and mesenchymal genes confirmed that cells treated with EZH2i express higher level of key mesenchymal genes at days 6 and 12 (Figure 4B). Therefore, we settled that inhibition of EZH2 enhances EMT and hinders MET in A549 cells.

To identify which are the genes directly regulated by EZH2 during EMT-MET we determined which promoters were bound by EZH2 and differentially regulated in cells treated with EZH2i. Out of the 829 reversibly regulated genes (identified in Figure 2D), we found that 335 genes (Clusters I+II+II+IV) were over-expressed on day 6 (Figure 4C, Table S1) or day 12 (Figure 4D, Table S1) in cells treated with EZH2i. Of these, 140 genes were marked by EZH2, H3K27me3 and H3K4me3 at their promoter region at day 0 (Figure 4E, F). These were very significantly enriched for EMT genes (Figure S4C) and included established regulators (i.e. *SNAI2*, *MMP2*, *MMP9*, *ITGB3* and *GREM1*) of different EMT processes (Table S2). In fitting with our previous analyses in Figure 3, binding of EZH2 to these genes was similar in epithelial and mesenchymal states, although their RNA expression was increased as cells acquired mesenchymal features on day 6 (Figure 4G, H). Importantly, inhibition of EZH2 did not hinder repression of epithelial genes at day 6 (Figure 4C), including *E-CADHERIN* (Figure S4D), indicating that EZH2 is not critical to downregulate expression of epithelial genes during EMT. RT-qPCR analysis confirmed that inhibition of EZH2 hindered repression of EZH2-target mesenchymal genes (*SNAI2* and *MMP2*), and led to expected secondary changes in the expression of *E-CADHERIN*, *N-CADHERIN* and *SNAI1* (Figure 4I, S4E). To address whether inhibition of EZH2 led to the acquisition of mesenchymal cellular features during EMT-MET, we measured the behaviour of EZH2i-treated cells in wound healing and soft agar assays. Inhibition of EZH2 led to increased healing ability and reduced capacity of anchorage-independent growth at days 6 and 12 (Figure 4J, K). Taken together, these experiments demonstrate that EZH2 represses a large set of mesenchymal genes during EMT-MET, and that blocking EZH2 methyltransferase activity favours the acquisition of molecular and cellular mesenchymal features.

**Figure 4.**
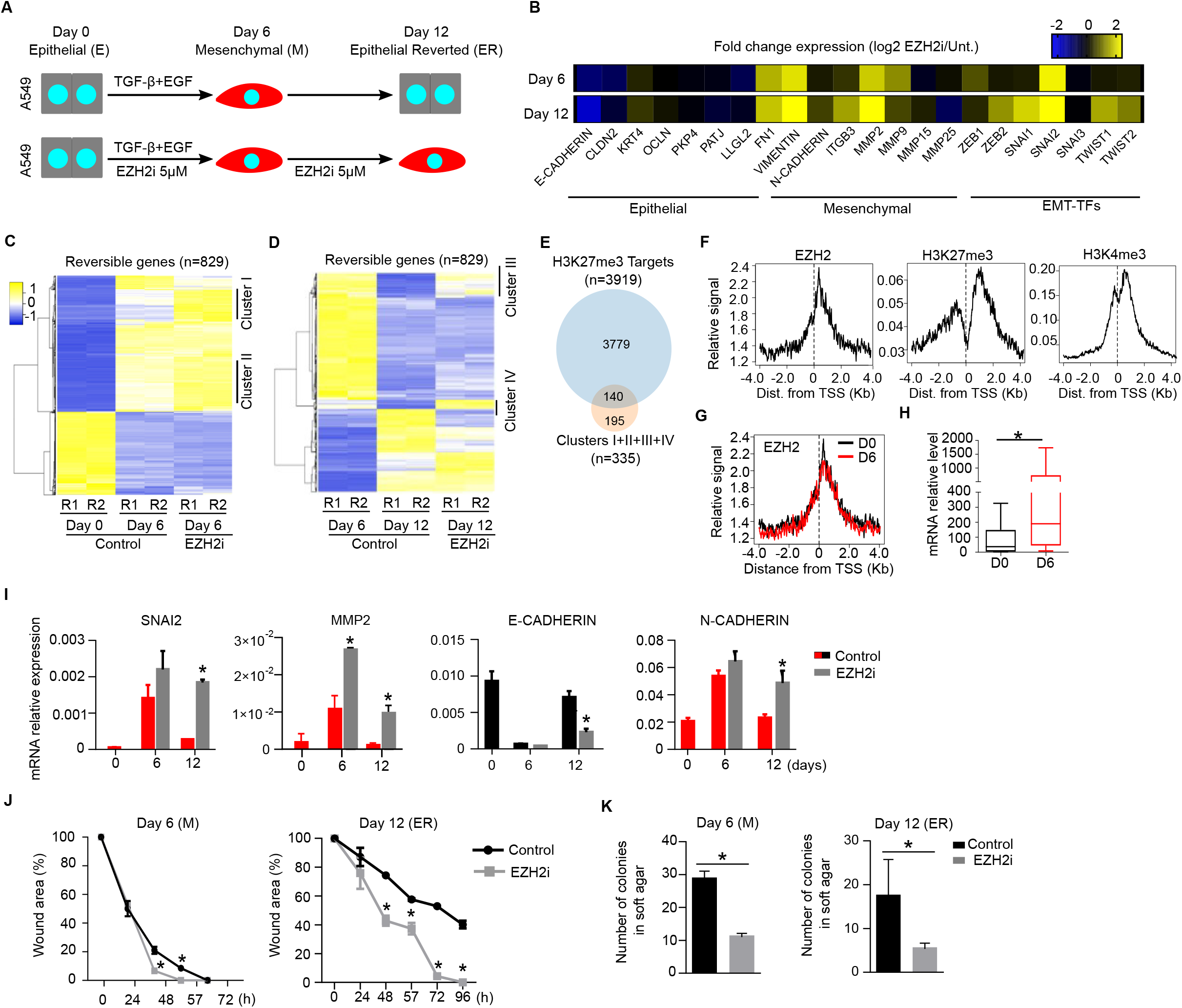
EZH2 represses a large set of mesenchymal genes during EMT-MET in A549 cells. **(A)** Schematic diagram of the experimental conditions used to study the effect of inhibiting EZH2 activity (EZH2i 5 μM) during reversible EMT-MET (TGF-β + EGF). **(B)** Heatmaps showing the fold change expression of epithelial and mesenchymal marker genes measured by cChIP-seq in cells treated with EZH2i vs untreated at day 6 and day 12. **(C, D)** Expression analysis of genes that are reversibly regulated (FC>2, p<0.05) during EMT-MET (n=829). Heatmaps show the expression of two biological duplicates (R1 and R2) at day 0 (E), day 6 (M) and day 12 (ER). Genes that are upregulated in EZH2i-treated cells are identified in cluster I-IV. **(E)** Venn diagram comparing H3K27me3 ChIP-seq signal at day 0 and genes in clusters I – IV. **(F)** Average binding plots of EZH2, H3K27me3 and H3K4me3 to the 140 overlapping genes identified in (E). **(G)** Average binding plots of EZH2 at day 0 and day 6 at the 140 overlapping genes identified in (E). **(H)** Box plot showing mRNA expression at day 0 and day 6 of the 140 overlapping genes identified in (E). **(I)** RT-qPCR analysis measuring mRNA level of indicated genes during EMT-MET in the absence or presence of EZH2i. Expression level is calculated relative to *GAPDH* and *ACTIN B*. **(J)** Graph comparing the kinetics of wound closure of cells at day 6 or day 12 of the EMT-MET treatment in the presence or absence of EZH2i. **(K)** Plot showing the number of colonies formed in soft agar by cells at day 6 or day 12 of the EMT-MET treatment in the presence or absence of EZH2i. Mean and SEM of 3 experiments are shown in I, J and K. Asterisks indicate statistical significance using a Mann-Whitney test (* p<0.05).

### EZH2 modulates the transcriptional upregulation of mesenchymal genes in response to TGF-β stimulation

To confirm that the phenotype observed in EZH2i-treated cells was a consequence of inhibiting EZH2 and not due to unspecific inhibition of another methyl transferase enzyme, we analysed the effect of another highly specific EZH2 inhibitor (EPZ6438) (Knutson et al. 2013). In agreement with previous results using the EZH2i GSK126, treatment of A549 cells with 2 μM EPZ6428 dramatically reduced the level of H3K27me3 on chromatin (Figure S5A) and promoted higher mRNA expression of EZH2 target genes (*SNAI2* and *MMP2*) during EMT-MET (Figure S5B). Concomitantly, cells treated with EPZ6428 displayed reduced expression of *E-CADHERIN* but increased number of *N-CADHERIN* transcripts (Figure S5B). Importantly, treatment with EPZ6428 also led to increased healing capacity in wound healing assays at both day 6 and 12 during EMT-MET assays (Figure S5C, D).

To obtain solid proof that EZH2 represses mesenchymal genes during EMT-MET in lung cancer cells, we used a genetic approach and derive A549 cells knock-out for EZH2 using CRISPR/Cas9 (Figure S5E). Upon transfection and flow cytometry sorting of the cells that had incorporated the GFP-expressing CRISPR/Cas9 plasmid, we derived both bulk and genetically clonal stable populations of A549 cells that expressed very low (bulk) or undetectable (clone) levels of EZH2 protein and H3K27me3 (Figure 5A). In fitting with the phenotype observed in experiments using EZH2 inhibitors, EZH2-null cells did not longer form tight epithelial colonies in culture, but organized as non-packed fusiform cells (Figure 5B). This was particularly evident in the clonal population of cells, suggesting that the bulk population harbours a low proportion of non-edited cells. In consonance with EZH2i-treated cells, EZH2-null cells displayed a proliferation capacity comparable to A549 parental cells (Figure 5C, S5F). The loss of EZH2 led to overexpression of EZH2-target genes *SNAI2* and *MMP2* coupled with changes in the expression of *E-CADHERIN* and *N-CADHERIN* (Figure 5D). In addition, loss of EZH2 repression induced acquisition of mesenchymal cellular features such as enhanced cell motility in wound healing assays, reduced anchorage-independent growth and decreased ability to form tumour spheres compared to parental cells (Figure 5E, S5G, H). Taken together, these experiments confirm that EZH2 activity is dispensable for cell growth but required to repress mesenchymal genes and uphold an epithelial identity in A549 lung cancer cells.

We wondered whether, given that EZH2-null cells already showed obvious mesenchymal molecular and cellular features, they could still respond to TGF-β stimulation and become more mesenchymal. We compared the expression of EZH2-target mesenchymal genes (*SNAI2*, *MMP2* and *ITGB3*) as well as *E-CADHERIN* and *N-CADHERIN* during *in vitro* EMT-MET in EZH2-null and parental cells. Non-stimulated EZH2-null cells already displayed a level of expression of mesenchymal genes similar to the one observed in TGF-β-treated parental cells on day 6 (see SNAI2 and MMP2 in Figure 5F). Interestingly, addition of TGF-β further enhanced transcriptional activity of EZH2-target genes SNAI2 and MMP2 (Figure 5F), and this was accompanied by the acquisition of an extreme fusiform morphology of EZH2-null cells at day 6 (Figure 5G). Importantly, removal of TGF-β from the culture media led to the restoration of the initial transcriptional level in both parental and EZH2-null cells for all genes tested (Figure 5F). Thus, we concluded that EZH2 binds and represses mesenchymal genes at different stages of the epithelial-mesenchymal spectrum where it seems to coordinate and buffer the transcriptional response of the cells to TGF-β stimulation (Figure 5H).

To determine whether the phenotype of EZH2 loss of function was orchestrated by SNAI2 protein we used CRISPR/Cas9 (Figure S5I) to derive a SNAI2-null bulk population of A549 cells. Inhibition of EZH2 activity induced the expression of SNAI2 protein in parental cells, but was barely detected in SNAI2-null cells (Figure S5J). SNAI2-null cells treated with EZH2i acquired a mesenchymal state similar to parental cells, as judged by their fusiform cell morphology as well as expression of *E-CADHERIN* and *N-CADHERIN* (Figure S5K, L). Importantly, in fitting with the role of SNAI2 as an inducer of cell motility ^26^, loss of function of SNAI2 blocked the acquisition of wound healing upon EZH2i-treament (Figure S5M). Thus, we interpreted that the mesenchymalization observed in cells with EZH2 loss of function depends on the combinatorial activity of several EZH2-target genes of which SNAI2 is responsible for changes in cell motility.

**Figure 5.**
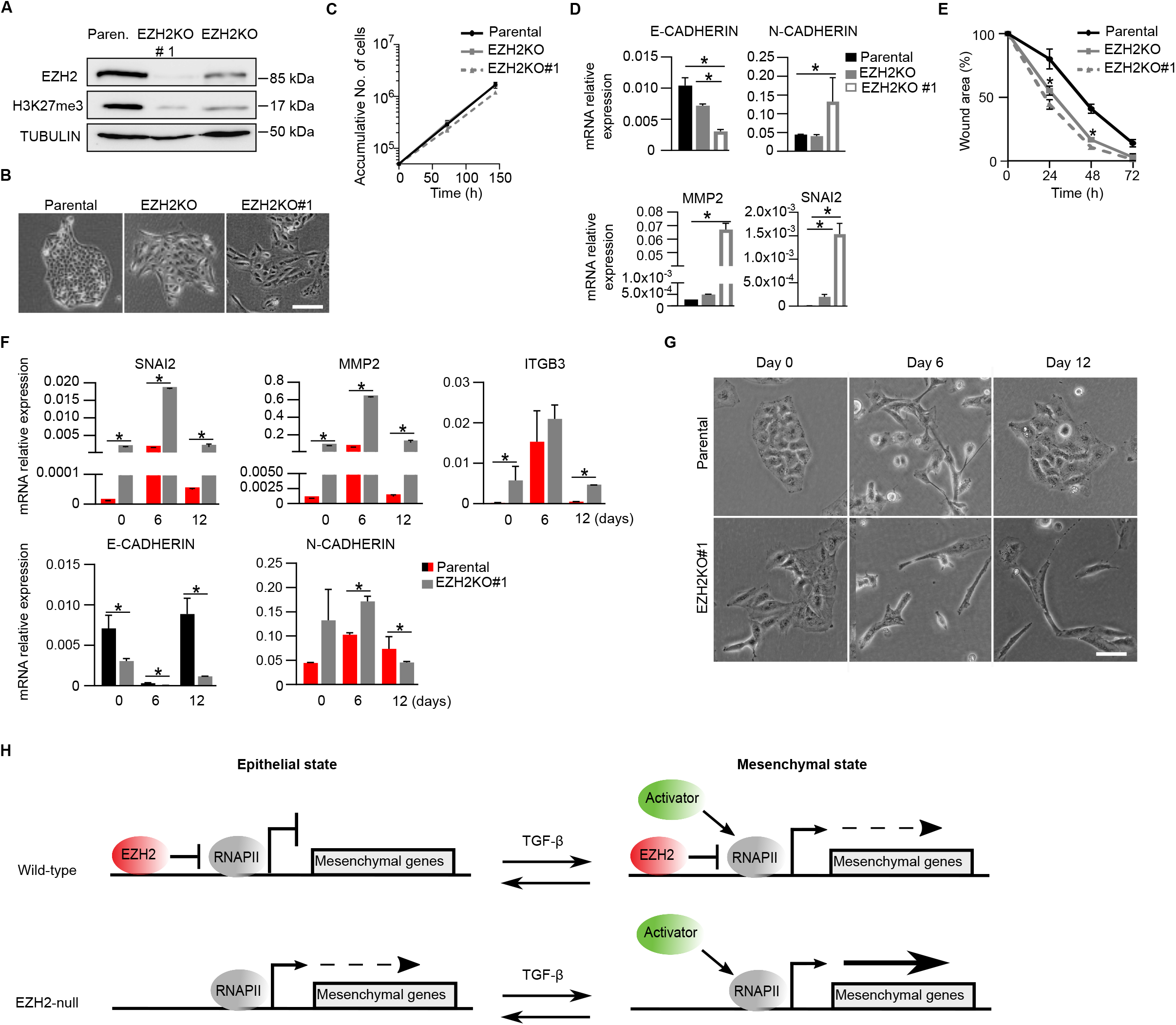
*EZH2*-null A549 cells acquire mesenchymal features but remain responsive to TGF-β stimulation. **(A)** Western blot analysis of whole cell extracts comparing the levels of EZH2 and H3K27me3 in parental and EZH2-null bulk (EZH2KO) and clonal (EZH2KO#1) populations. TUBULIN serves as loading control. **(B)** Brightfield images of parental, EZH2KO and EZH2KO#1 cells. Scale bar is 100 μm. **(C)** Growth curve of parental, EZH2KO and EZHKO#1 cells. **(D)** Analysis of mRNA expression by RT-qPCR in parental, EZH2KO and EZH2KO#1 cells. Expression level relative to GAPDH and ACTIN B is shown. **(E)** Kinetics of wound closure of parental, EZH2KO and EZH2KO#1 cells. **(F)** Analysis of mRNA expression by RT-qPCR in parental and EZH2KO#1 cells during EMT-MET. Expression level relative to GAPDH and ACTIN B is shown. **(G)** Brightfield images of parental and EZH2KO#1 cells during EMT-MET. Scale bar is 100 μm. **(H)** Schematic diagram proposing the mechanism by which EZH2 regulate mesenchymal genes during EMT-MET. EZH2, RNA Polymerase II (RNAPII) and a hypothetical transcriptional activator regulating mesenchymal genes in the presence or absence of TGF-β stimulation are depicted. Low or robust gene transcription is represented by discontinuous and solid arrows respectively. Mean and SEM of 3 experiments are shown in C, D, E and F. Asterisks indicate statistical significance using a Mann-Whitney test (* p<0.05).

### EZH2 protein level inversely correlates with expression of mesenchymal genes in human lung tumors

To obtain insights into whether EZH2 represses mesenchymal genes in human lung tumours, we analysed genome-wide datasets from the cancer genome atlas repository of protein and mRNA expression of resected tumours from 427 NSCLC patients. Expression of EZH2 protein showed a robust inverse correlation with the level of mRNA of mesenchymal genes identified in our *in vitro* system (Figure 6A). This correlation was solid for the set of H3K27me3-positive genes that were transcriptionally induced during EMT (386 genes identified in Figure 3F), and for the genes that were H3K27me3 positive and overexpressed in EZH2i-treated cells during EMT-MET (140 genes identified in Figure 4E). Inverse correlation was also evident for individual EZH2-targets including *SNAI2* and *MMP2* (Figure 6B). To further comprehend the role of EZH2 in lung cancer, we analysed whether the expression level of EZH2 was a marker of survival probability using available datasets of a cohort of 1925 NSCLC patients. In agreement with previous reports ^18,19,27^, expression of EZH2 mRNA was higher in lung tumours as compared to control healthy lung tissue (Figure S6A), supporting an oncogenic role of EZH2 in lung cancer. In fitting, high expression of EZH2 associated with poor survival (median survival EZH2, high: 45.27 months, low: 79.54 months) (Figure 6C). In contrast, other PRC2 subunits did not behave as markers of negative prognosis (Figure S6B), indicating that the effect of different PRC2 subunits in cancer progression is complex and must be individually analysed. We concluded that expression of EZH2 correlates with reduced expression of EZH2-target mesenchymal genes in lung cancer tumours and it is associated with poor survival.

**Figure 6.**
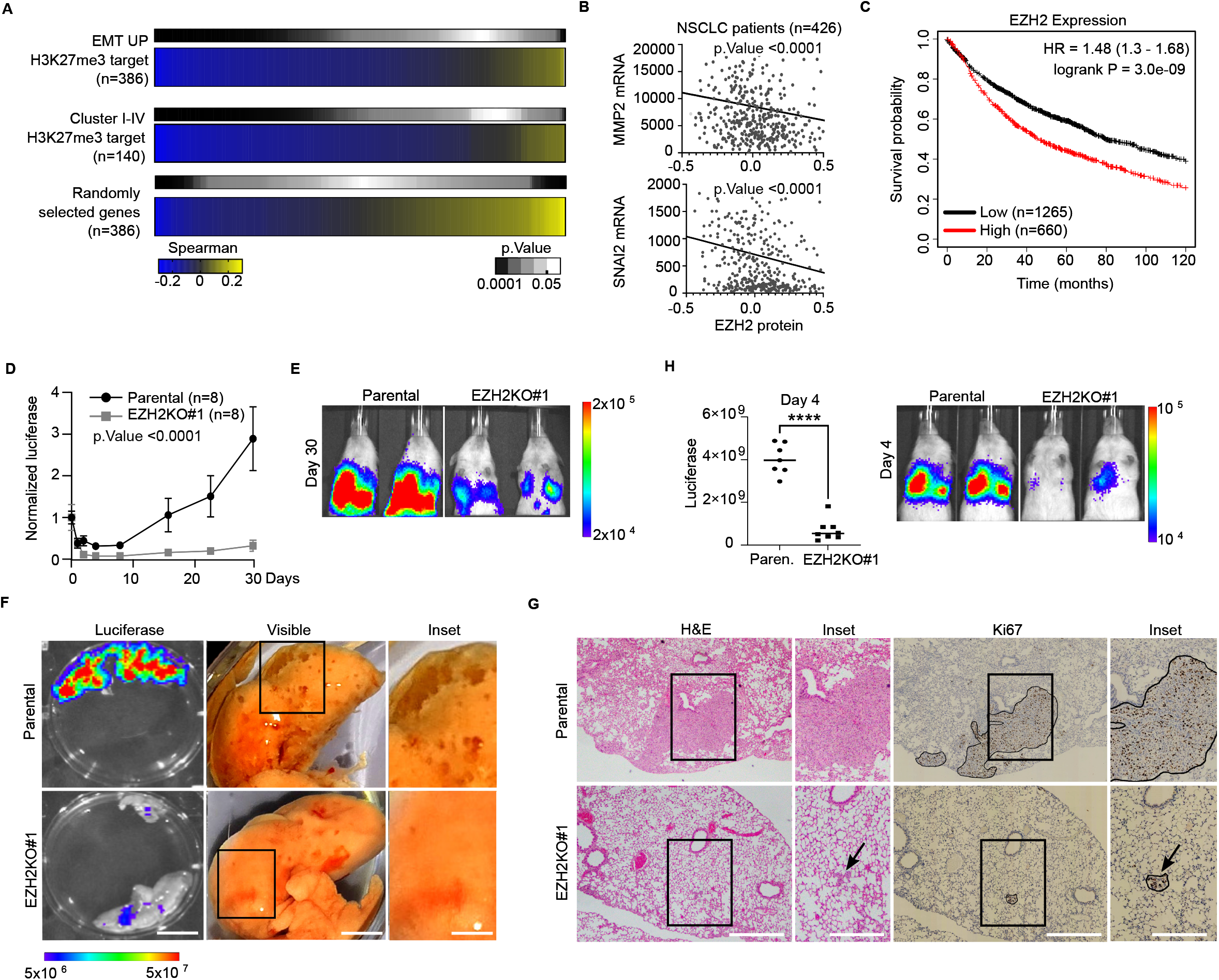
EZH2-null A549 cells display reduced tumor colonization capacity in xenograft assays. **(A)** Spearman’s correlation heatmaps of EZH2 protein and indicated sets of mRNAs in 426 human samples of NSCLC available at TCGA database. EMT UP and EZH2i responsive genes that are bound by H3K27me3 were identified in Figure 3F and Figure 4E respectively. The pattern of a set of randomly selected genes is shown for comparison purposes. **(B)** Spearman’s correlation of EZH2 protein and MMP2 (top panel) and SNAI2 (bottom panel) mRNA. Each dot represents one patient. **(C)** Kaplan-Meier plot showing survival probability of 1925 NSCLC patients depending on the expression of *EZH2* mRNA. **(D)** Kinetics of tumor growth inferred by luciferase activity upon tail intravenous injection of luciferase-expressing parental or EZH2KO#1 A549 cells in immunocompromised mice. Mean flux signal in eight mice was normalized to day 0 for every time point. Statistical significance was measure using an ANOVA test. **(E)** Representative bioluminescence images of animals at day 30 after injection. **(F)** Bioluminescence signal of surgically extracted mouse lungs 30 days post-injection (left panel). Scale bar is 10 mm. Image of lungs on day 40 post-injection. Scale bar is 5 mm. Insets highlight the presence of tumor-like nodules at the surface of the lungs. Scale bar is 2.5 mm. **(G)** Hematoxylin & eosin staining of fixed *ex vivo* lungs obtained from mice on day 30 after injection. Scale bar is 250 μm. Insets identify regions positive for H&E and Ki67 staining. Arrows highlight the position of Ki67-positive regions. Scale bar is 200 μm. **(H)** Histogram showing the bioluminescence signal of animals at day 4 after injection. Representative pictures are shown. Asterisks indicate statistical significance using a Mann-Whitney test (**** p<0.0001).

### EZH2-null A549 cells display reduced tumor colonization capacity in xenograft assays

Our analyses in cell lines and human patients support that EZH2 is required to transcriptionally repress the mesenchymal gene expression program during MET in lung cancer. Thus, we conjectured that EZH2 is required for tumour colonization and disease progression *in vivo.* To test this, we compared the tumour colonization capacity of A549 cells that were either wild-type or EZH2-null. Using lentiviruses, we transduced a luciferase reporter into EZH2KO#1 and parental cells, injected these luciferase-expressing cells intravenously in the tail vein of immunocompromised mice, and analysed tumour colonization kinetics by measuring luciferase activity *in vivo* for 30 days. Mice injected with A549 cells displayed robust luciferase signal in their thoracic cavity that increased during the course of the experiment (Figure 6D, E). In contrast, mice injected with EZH2-null cells showed very low luciferase activity that was sometimes indistinguishable from the background (Figure 6D, E). Examination of organs upon necropsy on day 30 showed that luciferase expression was evident in the lungs of mice injected with parental cells, while very low luciferase activity was detected in lungs from mice that had been injected with EZH2-null cells (Figure 6F, luciferase). No luciferase activity was found in brain nor liver in any of the two conditions (not shown). Although nodules on the surface of lungs were not evident on day 30, we could readily detect them 10 days later in mice injected with parental cells, but not in EZH2KO#1-injected mice (Figure 6F, visible). Haematoxylin-eosin and anti-Ki-67 staining of lung tissue samples confirmed the presence of tumour-like structures in histological preparations of parental xenografts, that were largely absent in mice injected with EZH2-null cells (Figure 6G). Importantly, close examination of the luciferase kinetics revealed that EZH2-null cells showed reduced colonization capacity on day 4 (Figure 6H), suggesting that reduced tumour formation capacity of EZH2-null cells stems from a failure to carry out MET during the homing process. Overall, we concluded that EZH2 is required to efficiently colonize lung tissue and form tumours in mouse xenograft models.

## Discussion

Metastasis is the cause of ~90% of cancer-associated mortality and current evidence strongly supports the idea that dynamic changes in cell state during epithelial–mesenchymal transitions are a critical part of the mechanism underlying metastatic dissemination ^7^. Therefore, characterizing the molecular basis of EMT and MET is crucial to better understand cancer progression and design more effective therapeutic approaches. During recent years studies have demonstrated the importance of signalling pathways and transcription factors in the regulation of EMT-MET. However, despite EMT-MET is essentially a change in gene expression that needs to be regulated at the chromatin level, little is known about which are the chromatin regulators involved as well as their mode of action ^3^. In this study, we have comprehensively analysed the role of the PRC2 catalytic subunit EZH2 in the regulation of EMT-MET in NSCLC. By combining epigenomics, cell biology and xenograft experiments, we show that EZH2 is required to repress mesenchymal genes during MET and for efficient tumour colonization. While these findings might be perceived as contradictory with initial studies showing that PRC2 facilitates EMT in other cancer systems ^21,22,28,29^, our observations are in fitting with more recent reports demonstrating that the PRC2 subunit EED can oppose EMT in lung cancer ^18^, and that the role of PRC2 in cancer is context-dependent ^14,30^. In this sense, it is important to note that PcG proteins repress hundreds of genes including proliferation inhibitors like *CDKN2A* that are frequently mutated in NSCLC patients and that regulate stress-response and entrance in S-phase through p14^ARF^ and p16^INK4A^ ^12,13^. Thus, in *CDKN2A* wild-type cancer cells, loss of function of PRC2 can reduce the fitness of the cells and indirectly affect cellular processes like EMT. Therefore, having used in this study human NSCLC cell lines that are *CDKN2A*-null was key to reveal that, on top of regulating cell proliferation through repression of *CDKN2A*, PRC2 maintains repression of mesenchymal genes and upholds epithelial identity in lung carcinoma cells. Moreover, having setup a reversible *in vitro* system that allowed us to interrogate the effect of inhibiting EZH2 during MET was instrumental to reveal the function of EZH2 as a repressor of the mesenchymal gene expression program in lung cancer. In fact, the absence of amenable systems to study the molecular basis of MET has probably hindered the discovery of chromatin regulators in this context. As a matter of fact, although studies have revealed chromatin modifiers (NuRD, G9a, SUV39H1/2) that silence epithelial genes during EMT ^4^, this is the first study to our knowledge that proposes that a chromatin regulator has a central role in the regulation of MET in cancer.

We found that EZH2 binds and represses mesenchymal genes that will be transcriptionally induced upon receival of EMT-inducing extracellular cues in lung carcinoma cells. This is reminiscent of the role of EZH2 in pluripotent embryonic stem cells (ESCs), where PRC2 binds and represses lineage specifier genes that will be activated when cell differentiation is triggered by extracellular signals ^9^. Loss of function of EZH2 in both systems results in upregulation of PRC2-target genes and associated changes in cellular features. However, while differentiation of ESCs involves dramatic changes in the genomic localization of PRC2 ^9,31^, is this study we showed that transition between the epithelial and mesenchymal states does not involve major changes in the genome-wide binding pattern of EZH2. In addition, we found that EZH2 remains functioning as a repressor of a similar set of genes at different stages of EMT-MET. Thus, the absence of a major genome-wide reorganization of EZH2 binding might be a defining epigenetic feature of EMT-MET. This is not surprising if we think about the nature of EMT-MET as compared to lineage transition during cell differentiation; while epithelial-mesenchyme transition requires metastable and reversible transition between cell states, cell differentiation is generally an irreversible process. Thus, constant binding of EZH2 to mesenchymal targets probably facilitates cell state reversibility, avoiding an irreversible change in cell identity. We propose that EZH2 promotes the residence of cancer cells within the epithelial-mesenchymal cell spectrum, endorsing them with cell plasticity and enhanced potential to adapt to new cellular niches during metastatic dissemination. This new facet of PRC2 activity in cancer is similar to the emerging function of PRC2 in postmitotic somatic cells, where it has been proposed to modulate gene expression in response to environmental cues ^32^.

The initial oncogenic role of EZH2 in cancers led to the development of several small molecule inhibitors ^19^, that are currently on clinical trials or even approved by FDA for the treatment of human cancer patients (i.e EPZ-6438 (tazemetostat) ^33^ in advanced epithelioid sarcoma). Importantly, later studies demonstrating that PcG proteins can have an oncogenic or tumour suppressive role, and that their activity depends on background mutations ^11,14^ is challenging the application of EZH2 inhibitors in the clinics. A direct presumption of the context-dependent function of EZH2 is that inhibiting EZH2 activity might be detrimental for a subset of patients, and therefore personalized treatments will be required to optimize the use of these drugs in clinical settings. Our findings suggest that treatment of patients with EZH2i will lead to different clinical outcomes depending on background mutations in *CDKN2A*. Treatment of patients with non-mutated *CDKN2A* cancer cells with EZH2i will lead to reduced cell proliferation, but will favour cell mesenchymalization and spreading of surviving cancer cells to other tissues of the body, where they might fail to efficiently colonize and form clinically-detectable metastasis. However, they might lead to micrometastasis that could produce harmful metastatic lesions later in life when treatment with EZH2i is discontinued. Importantly, if cancer cells are null for CDKN2A, treatment with EZH2i might not reduce cell proliferation but induce mobilization of cancer cells away from their residing tumours. In this case, inhibition of EZH2 will probably be detrimental for patients, specially at early stages of the disease. It is desirable that future studies challenge these predictions by combining mouse genetics, *in vivo* metastasis models and EZH2 inhibitors. In addition, we propose that stratification of cancer patients in clinical trials using EZH2 inhibitors will reveal key insights into the extent to which this dual role of EZH2 in proliferation and MET is important to optimize the treatment of human carcinomas.

## Methods

### Lung cancer cell lines and culture conditions

The A549 cell line was kindly provided by the lab of Dr. Maria Jose Serrano (GENYO, Granada, Spain). Cells were grown at 5% CO2 and 37 °C in DMEM high glucose media supplemented with 10% heat inactivated fetal bovine serum (FBS) (Gibco), penicillin/streptomycin (Gibco), L-glutamine (Gibco) and 2-mercaptoethanol (Gibco). H1944, H2122 and A427 cell lines were provided by the lab of Dr. Pedro P Medina (GENYO, Granada, Spain). These three cell lines were grown at 5% CO2 and 37° C in RPMI 1640 without L-Glutamine media supplemented with heat inactivated 10% FBS (Gibco), penicillin/streptomycin (Gibco), L- and 2-mercaptoethanol (Gibco). The four cancer cell lines are derived from human lung adenocarcinoma and harbor homozygous deletion of the *CDKN2A* locus and point mutations in *KRAS* locus resulting in a hyperactive form of the protein. Extended information about the genotype of the cell lines is provided in table S3.

### Induction of *in vitro* EMT-MET by TGF-β and EGF stimulation in A549 cells

Epithelial A549 cells were plated at a density of 10.000 cells/cm^2^ and treated with 10 ng/mL TGF-β (Prepotech) and 50 ng/mL EGF (Prepotech) during 6 days to induce transition into a mesenchymal state (EMT). Thereafter, both cytokines were removed from the culture media and cells were grown for 6 additional days to allow reversion to the epithelial state (MET). Cells were trypsinized, counted and replated at initial density in fresh media every 48h.

### Treatment of lung cancer cell lines with EZH2 inhibitors

EZH2 was inhibited using 5 μM GSK126 (A3446, APExBIO) or 2 μM EPZ-6438 (Tazemetostat) (S7128, Selleckchem). In treatments with EZH2 inhibitor for 12 days, cells were constantly maintained in the presence of GSK126 by refreshing it every two days as A549 cells were diluted to maintain exponential cell growth. During EMT-MET experiments, A549 cells were maintained in the presence of GSK126 or EPZ6438 during the course of the experiment. Inhibitors were refreshed every two days as cells were diluted to the corresponding density. In colony forming assays in the presence of EZH2 inhibitor, cells were pretreated with GSK126 for 4 days to reduce global H3K27me3 levels before plating the cells that were allowed to form colonies for 14 days in which the media was changed every 4 days refreshing GSK126. In growth curves, soft agar, tumor spheres and wound healing assays, cells were pretreated with GSK126 for 4 days to reduce global H3K27me3 levels before the start of the experiment. Cells were maintained in the presence of GSK126 during the course of the experiment as detailed below.

### Growth curve, colony formation and *in vitro* wound healing assays

For growth curves, cells were platted at density of 10.000 cells/cm^2^ and diluted before confluence to the initial density. Accumulative growth was calculated by applying the dilution factor used. Inhibition of EZH2 was carried out by pretreating cells with GSK126 for four days and refreshing the inhibitor in every cell dilution. Survival in the presence of cells treated with GSK126 was calculated relative to untreated cultures.

In colony formation assays, cells were plated at 100 cells/cm^2^ and maintained in culture for 14 days during which the media was refreshed every four days. Colonies were stained with 0.01% crystal violet (Sigma-Aldrich), counted and quantified with Image J.

For wound healing assays, A549 cells were grown to 100% confluence and a 1000 μL sterile pipette tip was used to produce a scratch in the monolayer of cells. Cells were grown in their culture media for 72-96 hour and the area of the wound is imaged every 24 hours and quantified using Image J software. The area of the wound at each point was normalized to the area of the wound at initial time.

### Formation of colonies in soft-agar and tumorspheres

Soft-agar colony formation was assayed as previously described ^34^ with modifications. A549 cells were embedded in 6-well culture plates at a density of 20.000 cells/well in 0.8% low melting point agarose (Thermo Scientific) on the top of a layer of low melting point agarose at 1.6%. 1 ml of DMEM was gently added into each well. Cells were then incubated for 21 days at 37°C and 5% CO2. The media was replaced every 4 days. When cells were treated with GSK126, they were pretreated for four days and the inhibitor was always added to the media. Colony formation was examined under a microscope after staining with 0.01% Crystal Violet (Sigma-Aldrich) and the colonies with a diameter over 100 μm were quantified using ImageJ.

Cells were seeded at a density of 1.000 cells/well in 24-well Costar Ultra-low Attachment Plates (Corning) in tumorsphere-forming media containing DMEM-F12 (Sigma-Aldrich), supplemented with 1X B27 (Thermofisher), 1 μg/ml hydrocortisone (Sigma-Aldrich), 4 ng/ml heparin (Sigma-Aldrich), 1X Insulin-Transferrin-Selenium (ITS -G) (Gibco), 1% penicillin/streptomycin (10.000 U/ml) (Gibco), 10 ng/ml fibroblast growth factor (FGF) (Sigma-Aldrich), 10 ng/ml hepatocyte growth factor (HGF) (Miltenyi Biotec), 10 ng/ml epidermal growth factor (EGF) (Sigma-Aldrich) and interleukin 6 (IL-6) (Miltenyi Biotec), as previously reported ^35,36^. Upon 72 hours of treatment, primary tumorspheres were harvested, trypsinized to single cells and re-plated in 24-well ultra-low attachment plates at a density of 1,000 cells/well in tumorsphere-forming medium to minimize clumping. Upon 72 hours, secondary tumorspheres with a diameter above 50 μm were quantified. Inhibition of EZH2 was achieved by pretreatment of cells with GSK126 during 4 days prior to the start of the experiment, and adding GSK126 to the tumor-sphere-forming media.

### Derivation of *EZH2* and *SNAI2* null cell lines by CRISPR/CAS9 in A549 cells

Single-guide RNA sequences for targeting *EZH2* and *SNAI2* genes were designed using Custom Alt-R CRISPR-Cas9 tool from Integrated DNA Technologies (IDT). The guide sequences were synthesized and cloned into the PX458 plasmid ^37^. A549 cells were transfected using lipofectamine 2000 (Thermofisher) using manufacturer’s recommended conditions. Cells positive for GFP expression were isolated using flow cytometry cell sorting and grown in culture to establish a bulk population of EZH2KO or *SNAI2*KO cells. Cells were maintained in culture for more than 30 days to confirm stability of the knockout. EZH2KO cells were plated at 1000 cells/cm^2^ to allow the formation of clonal colonies that were picked and expanded to establish clonal EZH2 knockout cells (EZH2KO#1). The sequences of the RNA guides are provided in supplementary table S3.

### RT-qPCR and Immunofluorescence

For RT-qPCR, RNA was isolated using Trizol reagent (Thermofisher), reverse transcribed using RevertAid Frist Strand cDNA systhesis kit (Thermofisher) and analyzed by SYBRG real-time PCR using GoTaq qPCR Master Mix (Promega). Primers used are provided in supplementary table S3.

Immunofluorescence analysis of A549 cells was carried out as described previously ^38^. Briefly, cells were fixed for 20 minutes in 2% paraformaldehyde, permeabilized 5 minutes in 0.4% Triton-X100 and blocked for 30 minutes in phosphate buffer saline supplemented with 0.05% Tween 20, 2.5% bovine serum albumin and 10% goat serum. Primary antibodies were Rb anti-EZH2 (CST) and Ms anti-H3K27me3 (Active Motif) were used. Secondary antibodies were goat anti-mouse alexa fluor 488 (Thermofisher), goat anti-rabbit alexa fluor 555 (Thermofisher). Slides were imaged by widefield fluorescence microscope Zeiss Axio Imager and images were analyzed using Image J. More information about antibodies used is provided in table S3.

### Histone extraction and Western blotting

Abcam protocol was used for acid extraction of histones cells. Briefly, cells were harvested, washed twice with ice cold PBS and resuspended triton extraction buffer (PBS with 0.5% TRITON X 100 (v/v), 2mM phenylmethylsulfonyl fluoride (PMSF), 0.02% (w/v) NaN_3_, 50 mM NaF, 10 μM Na_3_VO_4_ and protease inhibitor cocktail) at a density 10^7^ cells/ml. Cells were kept on ice for 10 minutes and then centrifuged at 6500 x g, 4°C for 10 mins. Nuclei were washed in half volume of TEB and centrifuged at 6500 x g, 4°C for 10 mins. Nuclei were resuspended in 0.2 N HCl at a density of 4×10^7^ nuclei/ml and incubate over night at 4°C. Samples were centrifuged at 6500 x g, 4°C for 10 mins. Histone-enriched supernatant was quantified using Bradford assay.

Western blots of whole cell extracts or histone preparations were carried out using standard procedures as previously described ^39^. The following primary antibodies were used: mouse anti-H3K27me3 (Active Motif), rabbit anti-Histone 3 (Abcam), rabbit anti-EZH2 (Diagenode), rabbit anti-SNAI1 (CST), rabbit anti-SNAI2 (CST), rabbit anti-ZEB1 (CST), mouse anti-E-CADHERIN (BD), mouse anti-ACTIN B (Sigma Aldrich), goat anti-LAMIN B (Santa Cruz), mouse anti-TUBULIN (Santa Cruz). Secondary species-specific antibodies conjugated to horseradish peroxidase were used: anti-rabbit-HRP (GE-Healthcare), anti-mouse-HRP (GE-Healthcare) and anti-goat-HRP (Abcam). Clarity Western ECL reagents (bio-rad) was used for detection. More information about antibodies used is provided in supplementary table S3.

### Calibrated chromatin immunoprecipitation sequencing (cChIP-seq)

EZH2 cChIP-seqs were carried out using a previously described protocol ^40^ with reported modifications ^41^ to quantify ChIP signal across different cell growth. Briefly, cells were trypsinized, resuspended in A549 cell-culture media, counted and 3.5×10^6^ A549 cells were mixed with 1.4×10^5^ mouse JM8 embryonic stem cells as spike-in control per reaction. Cells were spined and resuspended in 37°C complete media at a density of 5×10^6^ cells/ml and incubated in a rotating platform for 12 min with 1% formaldehyde at room temperature. After quenching formaldehyde fixation with glycine (125mM final concentration), cells were resuspended in swelling buffer (25 mM HEPES pH 7.9, 1.5 mM MgCl2, 10 mM KCl, 0.1% NP-40) at a density of 2×106 cells/ml. Nuclei were isolated using a dounce homogenizer (tight pestle; 50 strokes). Nuclei were resuspended in sonication buffer (1×107 cells/ml) and sonicated 1h full power 4°C (30sec ON /30sec Off). For immunoprecipitation, 1.25ug of EZH2 antibody (Diagenode) per one million cells was added to chromatin and incubated in a rotating wheel at 4°C overnight. Protein G magnetic beads (Dynabeads, Invitrogen) were added and chromatin was incubated for 5 hours. Washes were carried out for 5 min at 4°C with 1 ml of the following buffers: 1× sonication buffer, 1× wash buffer A [50 mM Hepes (pH 7.9), 500 mM NaCl, 1 mM EDTA, 1% Triton X-100, 0.1% Na-deoxycholate, and 0.1% SDS], 1× wash buffer B [20 mM tris (pH 8.0), 1 mM EDTA, 250 mM LiCl, 0.5% NP-40, and 0.5% Na-deoxycholate], and 2× TE buffer (pH 8). DNA was eluted in elution buffer (50 mM Tris pH 7.5, 1 mM EDTA, 1% SDS) and reversed cross-linked overnight at 65°C in 160mM NaCl and 20 ug/ml RNase, followed by 2h 45°C incubation with proteinase K (220 ug/ml).

For ChIP-qPCR analyses 1% of input was used. qPCRs were carried out by loading the same volume of DNA per sample and using GoTaq® qPCR Master Mix (Promega). Details of human-specific primers used are available in supplementary table S3.

For cChIP-seq, libraries of immunoprecipitated DNA were generated from 3 ng of starting DNA with the NEBNext Ultra DNA Library Prep kit for Illumina (#E7370) according to manufacturer’s instructions at the CRG Genomics Core Facility (Barcelona) and sequenced using a HiSeq 2500 Illumina technology. 20-30 million reads (50 bp single-reads) were obtained for each library.

Upon sequencing reads were aligned to the concatenated human and mouse reference genomes (hg19 and mm9) with Bowtie 2 ^42^ and the “–no-discordant’ and “–no-mixed” parameters. Multimapping reads and PCR duplicates were removed with SAMTools ^43^. To calibrate the samples, human reads were downsampled as previously ^41^. We used BamCompare from deepTools suite ^44^ to create bigwig files with the normalized signal against input sample. To identify EZH2-bound promoters, peak calling was performed with MACS2 ^45^ using input samples for background normalization and we considered as significant those peaks with a p<0.01. We annotated the peaks with HOMER ^46^ by defining promoter regions as +-2kb from Transcription Start Site (TSS) and using input samples for background normalization. Data mining of publicly available ChIP-seq datasets (H3K27me3 (GSM1003455) and H3K4me3 (GSM1003561) from ENCODE) were processed the same way. CoverageView (Coverage visualization package for R. R package version 1.20.0.) was used to calculate coverage around TSS. Average normalized reads for a genomic window of −0.5 kb to +1.5 kb relative to TSS was calculated for individual promoter regions and represented as boxplots. Average binding plots were generated by counting normalize reads every 10 bp.

### mRNA sequencing and data analysis

Analysis of mRNA expression in A549 cells during EMT-MET in absence or presence of GSK126 was performed in biological duplicates at days 0, 6 and 12. Total RNA was extracted using Trizol reagent (Thermofisher). Strand specific mRNA-seq libraries were generated using 500 ng of total RNA and the TruSeq stranded mRNA Library Prep kit (#20020595). Libraries were sequenced using HiSeq 2500 Illumina sequencer at the CRG Genomics Core Facility (Barcelona). 20-25 million (50 bp paired-end) reads were obtained for each sample. Raw fastq files were quality filtered and aligned to the Human genome (version hg19) using STAR 2.5.2 ^47^. Gene expression was normalized with trimmed mean of M values (TMM) ^48^ implemented in NOISeq R package ^49^. Differential expression analysis was performed with DESeq2 package ^50^. MA plot was generated with edgeR ^51^. Differentially expressed genes were defined as genes with that showed a log_2_ fold change greater than 1 or smaller than −1 with an adjusted p-value < 0.05. The list of reversible genes was the result from the intersection between the genes differentially expressed between day 6 and 0 and between day 12 and 6. Venny 2.1 online platform was used to create Venn diagrams. Gene ontology analyses were performed using “DAVID bioinformatics” and “ENRICHR”. The online tool Heatmapper and Graphpad Prism 9 were used to produce heatmaps graphs.

### *In vivo* xenograft bioluminescence assay

Transfection of packaging cells was carried using lipofection. Briefly, HEK293T cells were grown in DMEM (Cat#L0103-500, Biowest) supplemented with 10% fetal bovine serum (FBS), 100U/ml streptomycin and penicillin. Cells were plated on a 10-cm plates the day before transfection to ensure exponential growth and 80% confluence. pUltra-Chili-Luc vector plasmid (Addgene plasmid #48688), together with packaging (psPAX2, Addgene plasmids #12260) and envelope (pMD2.G, Addgene #12259) plasmids (18μg total DNA; plasmid proportions of 3:2:1, respectively) were resuspended in 0.5 ml free-serum DMEM and mixed at room temperature for 20 min with 45μL LipoD293 (Cat#SL100668, Signagen Laboratories, Rockville, MD, USA) previously diluted in 0.5 ml free-serum DMEM. The plasmid-lipoD293 mixture was added to cells. Transfection mixture was removed after 5h incubation and 7ml of total medium were added. The viral supernatants were collected at 48 and 72h, and filtered through a 0.45 mm filter (Cat#FPE404030, JET Biofil, Guangzhou, China), aliquoted and immediately frozen at −80°C. Viral titers (transduction units/ml) were calculated on the basis of the percentage of dTomato-positive cells detected in the linear range of a serial dilution of the supernatant. Viral titers were estimated in a highly permissive cell line such as K-562. Transduction of A549 cells and EZH2KO cells was carried out in three cycles on infection using 2mL of the supernatant containing lentiviral particles and 8 μg/mL polybrene in 6x multiwells. Cells with the same expression level of dTomato were sorted by flow cytometry and grown to found a stable population that expresses dTomato and luciferase.

Eight-week-old male NOD.scid Il2rg tm1Wjl/SzJ (NSG) mice (provided by the Animal Experimentation Unit at University of Granada) were intravenously injected in the vein of the tail with either EZH2KO#1 or parental A549 cell lines (1×10^6^ cells/mice) stably expressing Luciferase (two groups; eight mice per group). For bioluminescence imaging, mice were injected intraperitoneally D-luciferin dissolved in phosphate-buffered saline at a dose of 150 mg/kg body weight. After 5 to 8 minutes, the animals were anesthetized in the dark chamber using 3% isoflurane in air at 1.5 L/min and O_2_ at 0.2 L/min/mouse, and animals were imaged in a chamber connected to a camera (IVIS, Xenogen, Alameda, CA, USA). Exposure time was 3 min in *large binning*, and the quantification of light emission was performed in photons/second using Living Image software (Xenogen). Tumor growth was monitored at 0, 1, 2, 4, 8, 16, 23 and 30 days by *in vivo* imaging and bioluminescence measurement performed in ventral position. Between days 30 and 40, the two groups of mice were sacrificed, their lungs were extracted and their luminescence was measured *ex vivo*, and preserved in 10% formalin for immunohistochemistry analysis. After lungs were fixed in 10% formalin at 4 °C for 3 days, fixed tissues were embedded in paraffin and 4 mm sections were histologically processed for hematoxylin-eosin staining and immunohistochemistry using anti-human Ki67 antibody (DAKO #M7240, 1:75, 30 min at 37°C). All these procedures were performed using standardized protocols by Atrys Health S.A.

All procedures involving mice were performed in strict accordance with the recommendations in the Guide for the Care and Use of Laboratory Animals of the Bioethical Committee of the University of Granada and the protocol was approved by the Committee on the Ethics of Animal Experiments at the University of Granada. All procedures were performed under isoflurane inhalation anesthesia, and every effort was made to minimize suffering.

### Bioinformatic analysis of lung cancer patients

Correlation analyses of EZH2 protein and mRNA were carried out using C. Bioportal analysis tools and the TCGA-Firehose Legacy dataset containing data from 426 NSCLC patients. Kaplan meier survival analyses for human data were computed with the online tool Kaplan-Meier plotter and datasets containing 1925 NSCLC patients ^52^

### Statistical analyses

Statistical significance (p<0.05) was generally determined by applying a two-tailed non-parametric Mann-Whitney test. ANOVA test was used to determine significance of bioluminescence signal in mice injected with luciferase expressing cells. Spearman’s correlation coefficient was calculated to measure correlation among variables. All analyses were performed with graphpad prism 9.

## Data access

Datasets are available at GEO-NCBI with accession number GSE180067 (https://www.ncbi.nlm.nih.gov/geo/query/acc.cgi?acc=GSE180067) and will be released upon publication acceptance.

## Author contributions

DL designed and conceptualized the study. DL and AG designed experiments. AG, AM, HGA, LLO, JCAV, FEC and MEM performed and analysed experiments. JMM performed data analyses. PCS, SGP, LLO, PPM, ASP and DL provided scientific advice and resources. DL obtained funding and supervised research.

## Acknowledgments

We are very grateful to Diego de Miguel Perez and Dra. Maria Jose Serrano for providing cell lines and technical advice. We thank Dra. Aurora Serrano, Natasa Vukovic, core facilities at GENYO and the Animal Experimentation Unit at University of Granada for excellent technical support. We also thank the genomics unit at the CRG for assistance with RNA-seq and ChIP-seq experiments. This study was supported by the Spanish ministry of science and innovation (PID2019-108108-100, EUR2021-122005), the Andalusian regional government (PC-0246-2017, PIER-0211-2019, PY20_00681) and the University of Granada (A-BIO-6-UGR20) grants. Sergio Granados-Principal’s lab is funded by Instituto de Salud Carlos III (CP19/00029, PI15/00336, PI19/01533).

---

**Supplementary Table S1. List of genes identified in this manuscript in genome-wide analyses.**

**Supplementary Table S2. List of 140 genes directly regulated by EZH2 during EMT-MET.**

**Supplementary Table S3. Supplementary details of the reagents used in this study.**

## Figure legends

**Figure S1.**
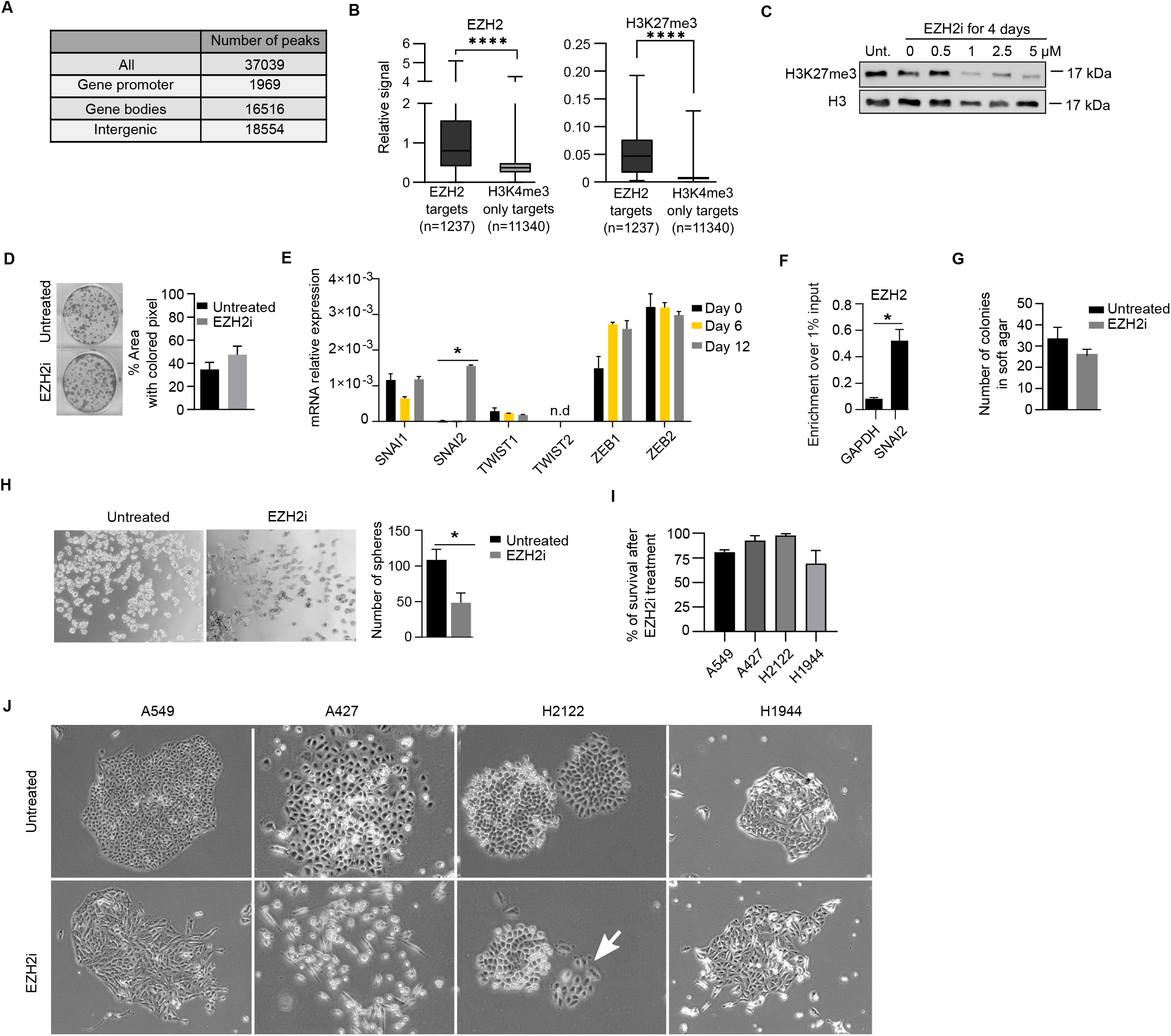
Inhibition of EZH2 leads to the acquisition of molecular and cellular mesenchymal features in NSCLC cells. **(A)** Table showing location of EZH2 binding peaks by cChIP-seq in A549 cells. **(B)** Boxplots of EZH2, H3K27me3 and H3K4me3 binding at the promoter regions (−0.5 to +1.5kb relative to TSS) of EZH2 target genes and control regions in A549 cells. **(C)** Western blot of histone extracts measuring the level of H3K27me3 in cells grown at indicated concentrations of EZH2i during 4 days in A549 cells. Total H3 total was used as loading control. **(D)** Representative images and quantification of crystal violet-stained colonies 18 days after plating at low density EZH2i-treated A549 cell cultures. **(E)** Analysis of mRNA expression by RT-qPCR of EMT-TFs in A549 cells treated with EZH2i. Expression level is calculated relative to *GAPDH* and *ACTIN B*. **(F)** Analysis of EZH2 binding by ChIP-qPCR at the promoter of SNAI2 in A549 cells. GAPDH promoter was used as a negative control. **(G)** Histogram showing the number of colonies formed after plating in soft agar EZH2i-treated A549 cells. **(H)** Tumor-sphere formation assay comparing untreated and EZH2i treated A549 cells. Representative images and quantitative histogram are shown. **(I)** Histogram showing cell survival upon EZH2i-treatment relative to untreated cultures for indicated NSCLC cell lines upon 4 days of growth. **(J)** Brightfield images of NSCLC cell lines plated at low density and 18 days of treatment with EZH2i. White arrow highlights a colony with mesenchymal morphology in H2122 cells. Mean and SEM of 3 experiments are shown in D, E, F, G, H and I. Asterisks indicate statistical significance using a Mann-Whitney test (* p<0.05, **** p<0.0001).

**Figure S2.**
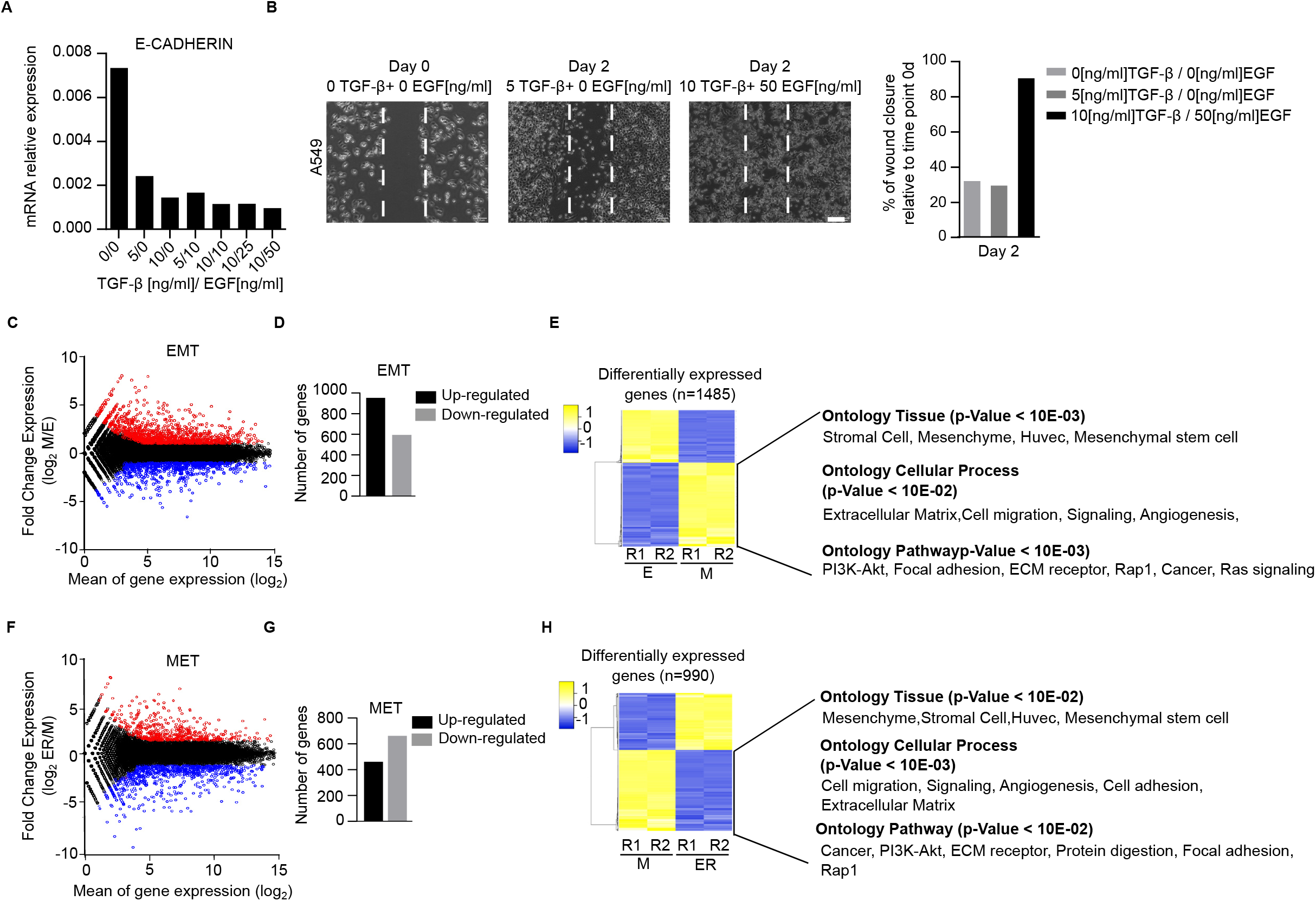
Analysis of global changes of gene expression during EMT-MET in A549 cells. **(A)** Histogram showing E-CADHERIN mRNA levels measured by RT-qPCR upon 48 hours of exposure to indicated concentrations of TGF-B and EGF. **(B)** Brightfield images and histogram quantifying wound healing capacity of cells that were treated with different concentrations of TGF-B and EGF. **(C)** MA plot analysis of mRNA datasets of cells during EMT: cells treated with 10 ng/ml TGF-B and 50 ng/ml EGF for six days (day 6, Mesenchymal, M) compared to untreated cells (day 0, Epithelial, E). Each dot represents one gene. Up-regulated genes are plotted in red and down-regulated genes in blue. **(D)** Histogram showing the number of significantly up-regulated and down-regulated genes during EMT. **(E)** Heatmap of significantly (FC>2, p<0.05) mis-regulated genes (n=1485) during EMT. Biological duplicates (R1 and R2) for day 0 (E) and day 6 (M) are shown. Gene ontology analyses of up-regulated genes is shown. **(F)** MA plot analysis of mRNA datasets of cells during MET: cells treated with 10 ng/ml TGF-B and 50 ng/ml EGF for six days (day 6, Mesenchymal, M) were allowed to revert to the epithelial state by removing TGF-B and EGF during the following six days (day 12, epithelial reverted, ER). Each dot represents one gene. Up-regulated genes are plotted in red and down-regulated genes in blue. **(G)** Histogram showing the number of significantly up-regulated and down-regulated genes during MET. **(H)** Heatmap of significantly (FC>2, p<0.05) mis-regulated genes (n=1485) during MET using. Biological duplicates (R1 and R2) for day 6 (M) and day 12 (ER) are shown. Gene ontology analyses of down-regulated genes is shown.

**Figure S3.**
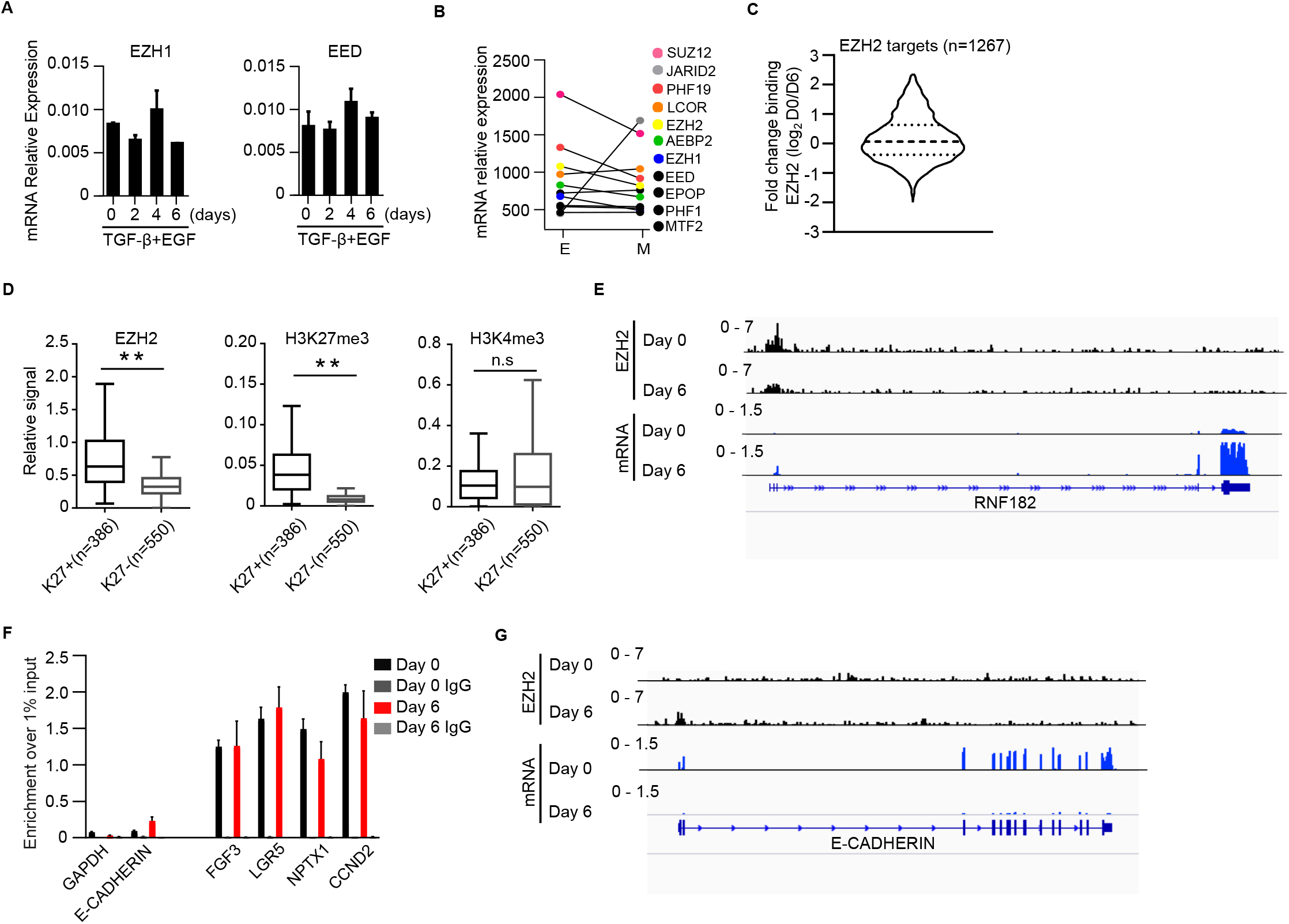
Low amount of EZH2 is recruited to *E-CADHERIN* promoter during EMT in A549 cells. **(A)** Expression of EZH1 and EED mRNA by RT-PCR at different times points during EMT. **(B)** mRNA expression of PRC2 subunits at day 0 (E) and day 6 (M) by RNA-seq. **(C)** Graph showing the fold change binding of EZH2 to target promoters (n=1267) at day 0 (E) and day 6 (M). **(D)** Box plot showing the binding of EZH2, H3K27me3 or H3K4me3 to EZH2-target promoters (n=1267) at day 0 (E). Asterisks indicate statistical significance using Mann-Whitney test (**p<0.01). n.s. means not statistically significant difference. **(E)** Genome browser view of EZH2 binding and mRNA expression at the *RNF182* locus at day 0 (E) and day 6 (M). **(F)** Binding of EZH2 to the promoter of EZH2-bound genes (*FGF3*, *LGR5, NPTX1, CCND2*) and *E-CADHERIN* at days 0 and 6 by ChIP-qPCR. *GAPDH* was used as a negative control. **(G)** Genome browser view of EZH2 binding and mRNA expression at the *E-CADHERIN* locus at day 0 (E) and day 6 (M).

**Figure S4.**
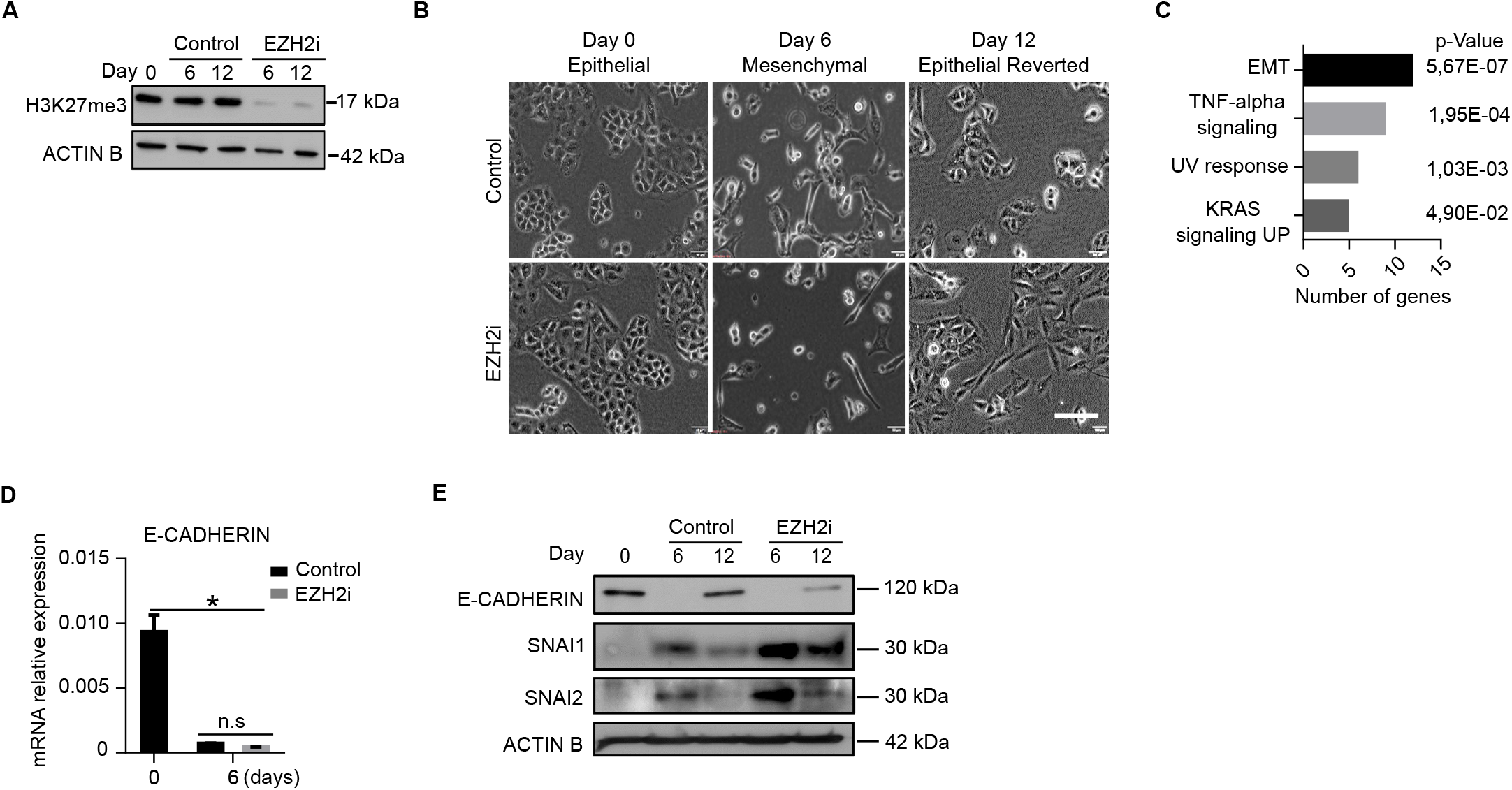
Inhibition of EZH2 does not hinder repression of *E-CADHERIN* during EMT in A549 cells. **(A)** Western blot analysis comparing levels of H3K27me3 in cells treated or untreated with EZH2i during EMT-MET. ACTIN B was used as loading control. **(B)** Brightfield images of cells at different stages of EMT-MET in the presence or absence of EZH2i. Scale bar is 100 μm. **(C)** Gene Ontology (GO) analysis using MSigDB Hallmark 2020 database of the 140 overlapping genes identified in Figure 4E. **(D)** Expression of E-CADHERIN mRNA by RT-qPCR during EMT in cells treated or untreated with EZH2i. Expression relative to *GAPDH* and *ACTIN B* is shown. Mean and SEM of 3 experiments is shown. Asterisks indicate statistical significance using a Mann-Whitney test (* p<0.05). **(F)** Western-blot of indicated proteins during EMT-MET carried out in the presence or absence of EZH2i. *ACTIN B* serves as loading control.

**Figure S5.**
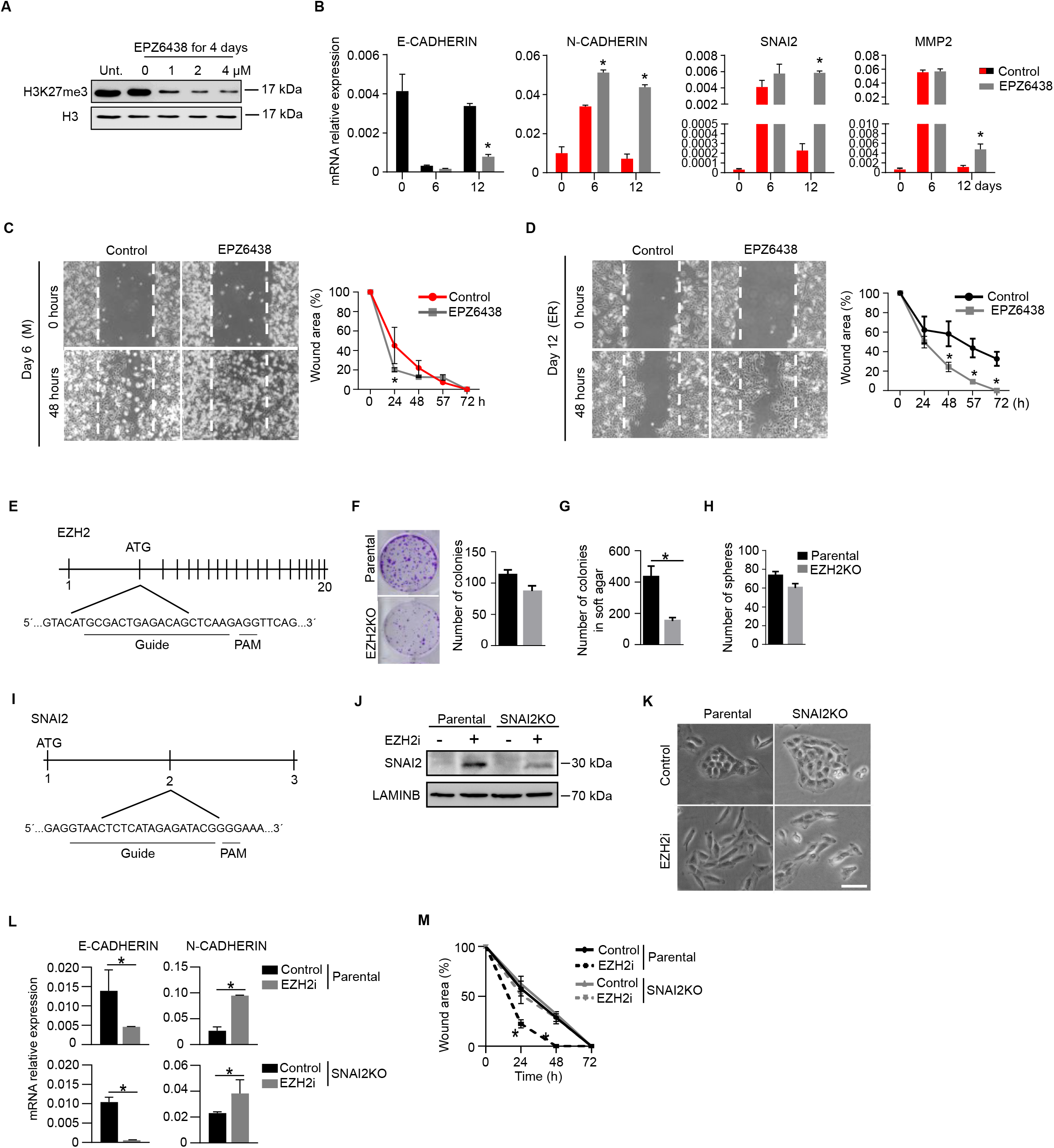
*SNAI2* is required for the acquisition of cell motility upon inhibition of EZH2 in A549 cells. **(A)** Western blot analysis of H3K27me3 level in histone extracts in presence of different concentrations of the EZH2 inhibitor EPZ6438 during 4 days. H3 was used as a loading control. **(B)** Analysis of mRNA by RT-qPCR during the course of EMT-MET in the presence or absence of 2 μM EPZ6438. Expression relative to GAPDH and ACTIN B is shown. **(C, D)** Brightfield images and graph of a wound healing assay comparing wound closure ability of cells at day 6 (C) or day 12 **(D)** of EMT-MET in the presence or absence of 2 μM EPZ6438. **(E)** Scheme showing the CRISPR/CAS9 strategy used to knock out EZH2 in A549 cells. Exons are depicted as black bars. Location of the ATG as well as the exon targeted by the guide RNA and the PAM sequence are indicated. **(F)** Images and histogram of a colony formation assay 18 days after plating at low density. **(G)** Histogram showing the number of colonies after 18 days of growth in soft agar by parental and EZH2KO cells. **(H)** Histogram showing the number of tumoral spheres produced by parental and EZH2KO cells. **(I)** Scheme showing the CRISPR/CAS9 strategy used to knock out SNAI2 in A549 cells. Exons are depicted as black bars. Location of the ATG as well as the exon targeted by the guide RNA and the PAM sequence are indicated. **(J)** Western blot analysis of whole cell extracts of parental and SNAI2KO A549 cells after 12 days of treatment with EZH2i. LAMINB was used as a loading control. **(K)** Brightfield images of parental and SNAI2KO treated for 12 days or not with EZH2i. **(L)** Analysis of mRNA expression by RT-qPCR in parental and SNAI2KO cells after 12 days of treatment with EZH2i. Expression level relative to GAPDH and ACTINB is shown. **(M)** Plot analyzing the kinetics of wound closure of SNAI2KO and parental cells upon 12 days of treatment with EZH2i. Mean and SEM of 3 experiments are shown in B, C, D, F, G, H, L and M. Asterisks indicate statistical significance using a Mann-Whitney test (* p<0.05).

**Figure S6.**
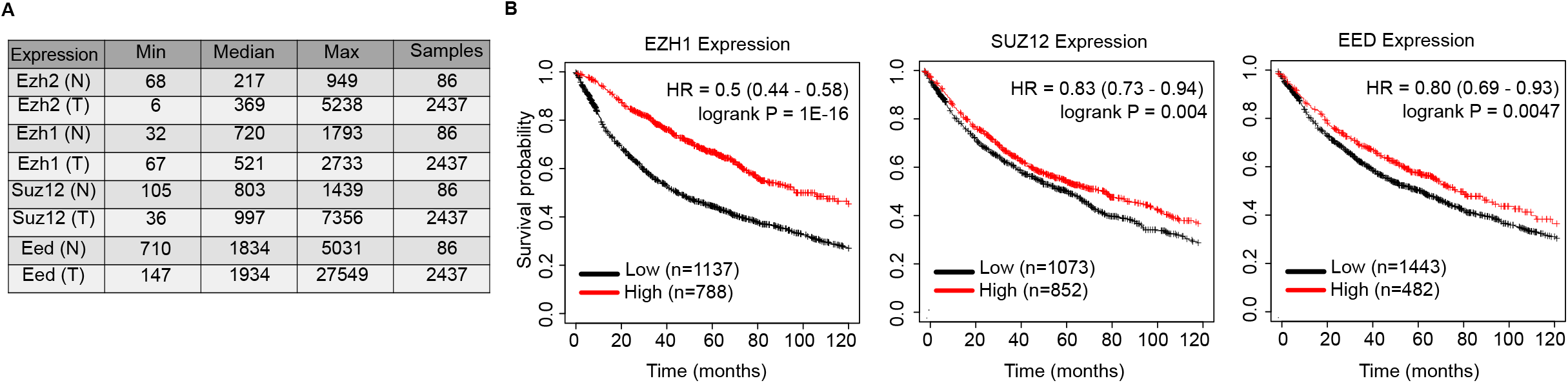
EZH2 is overexpressed in lung tumors. **(A)** Table showing the minimum, median and maximum expression values of several PRC2 subunits mRNA in samples from normal (N) (n=86) and tumoral (T) (n=2437) lung tissue. **(B)** Kaplan-Meier curve showing survival probability of 1925 NSCLC patients depending on the expression of *EZH2, SUZ12 or EED* mRNA.

