## Supplemental Table S2 for "EZH2 endorses cell plasticity to carcinoma cells facilitating mesenchymal to epithelial transition and tumour colonization"

**Genes falling within clusters I, II, III and IV and positive for H3K27me3 (n=140) (Figure 4E)**

| Gene | Process |
|---|---|
| ADORA1 | Cell surface |
| ADRA2A | Cell surface |
| ANTXR2 | Cell surface |
| CD200 | Cell surface |
| CD82 | Cell surface |
| CDCP1 | Cell surface |
| CDH11 | Cell surface |
| CHRNA3 | Cell surface |
| CNIH3 | Cell surface |
| DNER | Cell surface |
| EPHA4 | Cell surface |
| EPHB1 | Cell surface |
| EPHB6 | Cell surface |
| FAT3 | Cell surface |
| FXYD5 | Cell surface |
| GABRQ | Cell surface |
| GJB2 | Cell surface |
| GNG4 | Cell surface |
| GRIN3B | Cell surface |
| GXYLT2 | Cell surface |
| HRH1 | Cell surface |
| HTR7 | Cell surface |
| KCNA7 | Cell surface |
| KCNC3 | Cell surface |
| KCNMA1 | Cell surface |
| KCTD16 | Cell surface |
| NTSR1 | Cell surface |
| PAEP | Cell surface |
| PCDH1 | Cell surface |
| PODXL | Cell surface |
| PROKR1 | Cell surface |
| PTPRN | Cell surface |
| RASGRP1 | Cell surface |
| RASGRP3 | Cell surface |
| RET | Cell surface |
| SERPINE2 | Cell surface |
| SLAMF9 | Cell surface |
| SLC2A13 | Cell surface |
| SLC6A15 | Cell surface |
| SLC7A8 | Cell surface |
| SLCO5A1 | Cell surface |
| SRPX | Cell surface |
| TMEM255E | Cell surface |
| TSPAN5 | Cell surface |
| VSTM4 | Cell surface |
| AMOT | Cytoskeleton |
| ARC | Cytoskeleton |
| CORO2B | Cytoskeleton |
| DAAM1 | Cytoskeleton |
| FHOD3 | Cytoskeleton |

| Gene | Process |
|---|---|
| FMNL3 | Cytoskeleton |
| KANK4 | Cytoskeleton |
| KY | Cytoskeleton |
| LARP6 | Cytoskeleton |
| MYO1D | Cytoskeleton |
| NAV1 | Cytoskeleton |
| PRR5L | Cytoskeleton |
| STMN3 | Cytoskeleton |
| ADAM19 | Extracellular Matrix |
| APOLD1 | Extracellular Matrix |
| COL13A1 | Extracellular Matrix |
| COL5A1 | Extracellular Matrix |
| CPE | Extracellular Matrix |
| CTHRC1 | Extracellular Matrix |
| FBN1 | Extracellular Matrix |
| FURIN | Extracellular Matrix |
| ISM2 | Extracellular Matrix |
| ITGA8 | Extracellular Matrix |
| ITGB3 | Extracellular Matrix |
| MMP2 | Extracellular Matrix |
| MMP9 | Extracellular Matrix |
| STC1 | Extracellular Matrix |
| TIMP3 | Extracellular Matrix |
| BMP2 | Ligand |
| BMP6 | Ligand |
| CCL7 | Ligand |
| CRLF1 | Ligand |
| CSF1R | Ligand |
| FGF1 | Ligand |
| GAL | Ligand |
| GDF6 | Ligand |
| GREM1 | Ligand |
| IGFBP5 | Ligand |
| IGSF9 | Ligand |
| INHBA | Ligand |
| NKD1 | Ligand |
| SEMA3E | Ligand |
| SEMA7A | Ligand |
| TNFAIP6 | Ligand |
| VGF | Ligand |
| WNT5B | Ligand |
| WNT9A | Ligand |
| BTBD11 | Other |
| CHST6 | Other |
| COLEC12 | Other |
| CYP26B1 | Other |
| EGLN3 | Other |
| GFPT2 | Other |
| HK2 | Other |
| HS3ST3B1 | Other |

| Gene | Process |
|---|---|
| LIPG | Other |
| NOVA2 | Other |
| OXCT1 | Other |
| PLD5 | Other |
| PLPPR5 | Other |
| SUSD4 | Other |
| SYT11 | Other |
| UBTD1 | Other |
| WSCD1 | Other |
| XYLT1 | Other |
| BEAN1 | Other |
| CDYL2 | Regulation of transcription |
| CREB3L1 | Regulation of transcription |
| EGR1 | Regulation of transcription |
| EGR2 | Regulation of transcription |
| FOSB | Regulation of transcription |
| FOXP1 | Regulation of transcription |
| INSYN2B | Regulation of transcription |
| MAF | Regulation of transcription |
| NR4A2 | Regulation of transcription |
| NRIP3 | Regulation of transcription |
| SNAI2 | Regulation of transcription |
| TSHZ3 | Regulation of transcription |
| VDR | Regulation of transcription |
| ZNF703 | Regulation of transcription |
| DACT1 | Signaling |
| FHL1 | Signaling |
| GRB10 | Signaling |
| HHIP | Signaling |
| INPP4B | Signaling |
| KIAA1549L | Signaling |
| MOB3B | Signaling |
| PDE11A | Signaling |
| PIK3AP1 | Signaling |
| PIK3CD | Signaling |
| PKIA | Signaling |
| PPP1R14C | Signaling |
| SLN | Signaling |
| SPHK1 | Signaling |
| TNFAIP8L3 | Signaling |
